# OrthoGLMM: Phylogenetic Association Testing for Gene Content and Trait Evolution

**DOI:** 10.64898/2026.07.03.736217

**Authors:** Joseph Guhlin, Phoebe Keddell, Peter Dearden

## Abstract

1

**Motivation:** Comparative genome projects can now assemble and annotate hundreds of species, creating an opportunity to test whether species-level traits are associated with repeated changes in gene content. These tests must account for shared ancestry, sparse orthogroups, rare trait origins, and thousands of simultaneous associations.

**Results:** We present OrthoGLMM, a phylogenetically informed framework for the association of traits and orthogroup presence/absence or copy number across species. OrthoGLMM combines deterministic GLMM scans with solver-rerun empirical calibration and calibrated FDR estimation. In three benchmark datasets, OrthoGLMM recovered expected signals for bacterial diazotrophy, plant nodulation, and marine mammals.

**Availability and Implementation:** Source code, documentation, example data, and reproducibility scripts will be available at http://github.com/jguhlin/OrthoGLMM.

## 2 Introduction

Understanding how traits evolve and are lost is a central goal of comparative genomics and evolutionary biology. Traits can arise independently in related lineages^1–4^, be inherited from a common ancestor, or be lost after evolving^5,6^. Molecular routes to trait evolution include sequence-level change^1,7^, regulatory change^8–12^, and gene-content variation^13–16^. Gene-content variation includes gene gain and loss, horizontal gene transfer, copy-number change, and gene-family expansion or contraction. These patterns motivate the concept of genetic toolkits: shared gene sets repeatedly recruited during parallel evolution of similar traits^17–20^.

Repeated trait evolution can leave molecular signals that are comparable across lineages. In bacteria and archaea, nitrogen fixation reflects the evolutionary history of nitrogenase genes and associated cofactor-biosynthesis machinery, including gene gain, loss, duplication, and horizontal transfer^21^. In plants, C4 and CAM photosynthesis have each evolved multiple times from C3 ancestors, with independent C4 origins in grasses repeatedly recruiting members of the same gene families^22–25^. Similar comparative patterns have been described for fungal hyphae and multi-cellularity, eusociality in insects, mammalian echolocation, and electric organs in fishes, where repeated phenotypic evolution can involve conserved molecular machinery, parallel recruitment of paralogs, lineage-specific gene gain and loss, or convergent regulatory change^7,26–29^. Gene-content evolution also includes loss and contraction; for example, marine mammals often show reduced chemoreceptor repertoires^30^. These examples show that similar traits can be associated with repeated changes in genes, gene families, and gene repertoires across the tree of life.

Because species share ancestry, gene presence/absence and copy-number variation can co-occur through inheritance from a common ancestor. Downstream analyses therefore need to control for phylogenetic signal^5,31–33^. Comparative genomics is now sufficiently data-rich for high-throughput genome-content screens. Genome assembly is increasingly inexpensive and fast for many taxa^34–38^. Public genomes can be assembled or annotated at scale39,40, and multiple tools assign genes into orthologous groups^41–44^, including internal node-based hierarchical orthogroups (HOGs)^45^.

In population genomics, genome-wide association studies (GWAS) test genetic variants against phenotypes and have identified trait-associated loci across many systems^46,47^. There are multiple examples of experimental validation of candidate genes^48–51^, and some genes recur in independent analyses across related species^52^. GWAS frameworks are primarily designed for population-level variation anchored to a reference genome, such as single-nucleotide polymorphisms (SNPs), with relatedness modeled among individuals. Phylogenetic regression methods provide tools for testing associations among species-level traits while accounting for shared ancestry; packages such as PhyloGLM^53^, MCMCglmm^54^, and brms^55^ fit targeted phylogenetic generalized or Bayesian mixed models. Adjacent comparative tools address gene-family evolution, binary-trait association on phylogenies, or phylogeny-aware genotype-phenotype association, including CAFE5^56^, SimPhyNI^57^, CALANGO^58^, and Evolink^59^. Comparative genome-content screens still pose a distinct inference problem: tens of thousands of orthogroups vary in prevalence, copy-number distribution, sparsity, and clade structure, while the focal trait may have few independent origins.

We present OrthoGLMM, a package for high-throughput, phylogenetically informed, hypothesis-generating scans of orthogroup presence/absence and copy number. OrthoGLMM combines deterministic association testing with solver-rerun empirical calibration and calibrated FDR estimation. Tests can be run using a species tree or phylogenetic covariance matrix, with separate modes for orthogroup presence/absence and copy number. Optional follow-up layers group related orthogroups into meta-families or prioritize candidate orthogroups for downstream analyses of xenology, sequence evolution, or molecular convergence using tools such as HyPhy^60^, BUSTED-PH^61^, PAML^62^, and TraitRelax^63^.

We test OrthoGLMM on real datasets with well-studied traits. Specifically, we test associations between orthogroups and diazotrophy in 400 bacterial genomes, nodulation-related traits in 89 plant genomes, and marine lifestyle in 106 mammalian genomes.

## 3 System and methods

A selection of genomes was downloaded from NCBI^37^ between 28 January and 4 February 2026, comprising 400 bacterial, 89 plant, and 106 mammalian genomes. Traits were identified from curated phenotypic resources: bacterial diazotrophy from FAPROTAX genus-level annotations^64^, plant nodulation from the Afkhami et al. Dryad dataset^65^ with manually curated non-nodulating outgroups, and mammalian marine status from the Society for Marine Mammalogy species list^66^ with a manually curated terrestrial comparison set. Traits were encoded as 1 for positive states: diazotrophy, nodulation, or marine lifestyle.

Bacterial genomes were annotated with Prodigal v2.6.3^67^, and eukaryotic genomes were annotated with Galba v1.0.11^39^. This was done to reduce annotation-method bias^68^. Orthogroups were created using OrthoFinder v3.1.2^41^. BUSCO v6.0.0 was used to assess gene annotation completeness with the ODB10.1 bacteria_odb10, embryophyta_odb10, and mammalia_odb10 lineages^69^.

Putative functions for top-ranking genes were assigned with DIAMOND v2.1.21^70^ blastp against the UniProt/Swiss-Prot database^71^, downloaded on 29 December 2025. Representative sequences were taken from OrthoFinder outputs.

## 4 Algorithm

OrthoGLMM takes as input an orthogroup count or feature matrix, a species phylogeny or phylogenetic covariance matrix, and trait values (Figure 1). Optional follow-up analyses can also use orthogroup gene trees. For the exemplar analyses, OrthoFinder supplied orthogroups, resolved gene trees, and a STAG species tree^41,72^.

**Figure 1.**
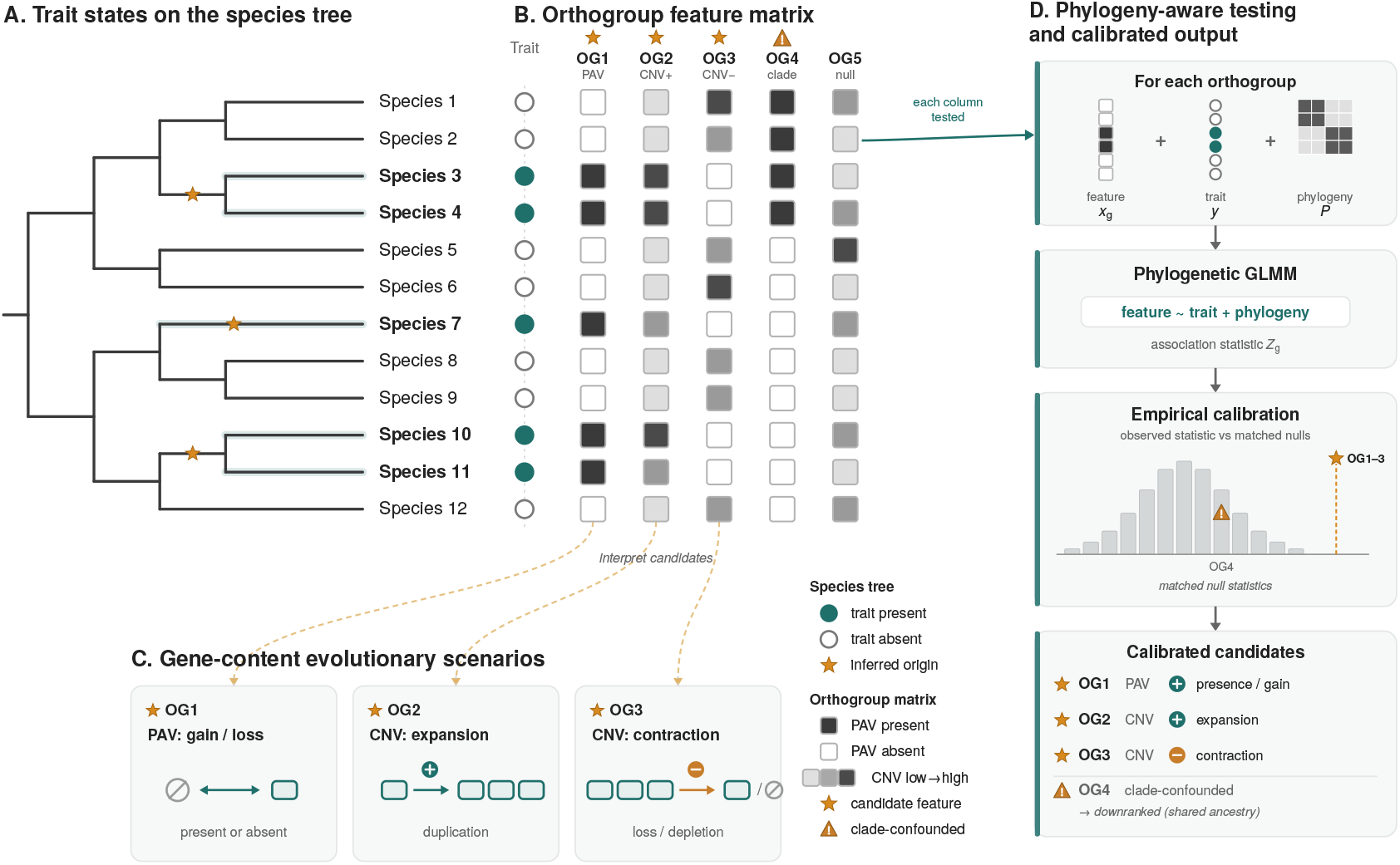
Conceptual overview of OrthoGLMM: from phylogenetically structured gene content to calibrated candidate gene families. **(A)** Rooted species tree of 12 taxa with trait states at the tips (filled circle, trait present; open circle, trait absent); stars (★) mark putative independent origins of the trait. **(B)** Orthogroup feature matrix in the same species order, scored as presence/ absence (PAV; filled, present; open, absent) or copy number (CNV; light→dark, low→high). Stars (★) mark candidate features (OG1 presence/absence, OG2 expansion, OG3 contraction); the warning symbol 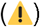 marks a clade-confounded feature (OG4); OG5 is background variation. **(C)** Interpretation of the candidate columns as gene-content histories: gain/loss, expansion, or contraction/depletion. **(D)** For each orthogroup, the feature (*x*_*g*_), trait vector (*y*), and phylogenetic covariance matrix (*P*) enter a phylogenetic GLMM (feature trait + phylogeny), giving the association statistic 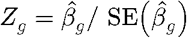. Observed statistics are compared against matched empirical null distributions from solver-rerun calibration: tail features are retained (★) while the clade-confounded feature lies within the null and is downranked 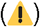. Calibrated p-values and Benjamini–Hochberg FDR yield a ranked candidate list rather than raw matrix pattern matches.

During preprocessing, OrthoGLMM checks input dimensionality, species order, covariance validity, finite entries, and required metadata. Matrices are aligned to the same species order, and entries with missing phenotype values are dropped. Covariance matrices are symmetrized on loading; non-finite entries raise an error, and non-positive-definite matrices produce explicit warnings before solver-level stabilization.

Given a species tree, OrthoGLMM creates a phylogenetic relatedness matrix *P*. OrthoGLMM uses the root encoded in the input Newick tree, so unrooted trees must be rooted before analysis. Non-ultrametric trees are ultrametricized before matrix construction. For each pair of species, covariance is defined as the shared path length from the root to their most recent common ancestor:

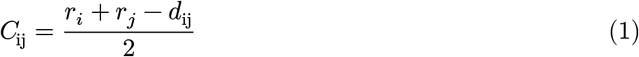

Here, *r*_*i*_ and *r*_*j*_ are root-to-tip distances, and *d*_ij_ is the patristic distance between species. Small negative eigenvalues are shifted before standardization. The covariance matrix is standardized to a correlation matrix with unit diagonal and used as a phylogenetic covariance term.

OrthoGLMM tests each orthogroup in two modes: presence/absence and copy number. Counts are converted either to a binary indicator of orthogroup presence or to a rank-inverse-normalized copy-number response. Presence/absence is fit with a Bernoulli/logit mixed model, whereas normalized copy number is fit with a Gaussian mixed model. The orthogroup-specific trait effect is used as the association statistic, with final significance assessed after empirical calibration and FDR correction.

For each orthogroup, the raw test statistic is the estimated trait effect divided by its standard error:

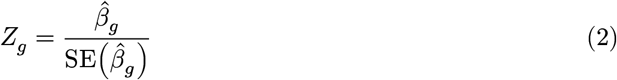

Raw two-sided Wald p-values are diagnostic screening values. Final inference uses empirical solver-rerun calibration. OrthoGLMM generates matched null inputs, reruns the same solver with the same feature mode, covariates, backend, and tuning settings, and compares each observed statistic with the resulting empirical null distribution.

The pipeline supports auto null-scheme selection: with a supplied species tree it selects `fixed_trait_clade`, and otherwise it selects `phylo_matched`. For singular or near-singular trait origins, `fixed_trait_clade` holds the observed phenotype and covariates fixed while shuffling orthogroup response vectors within tree-defined clade blocks. For convergent gain or loss traits, `phylo_matched` draws matched null phenotypes from the phylogenetic covariance while preserving binary prevalence or the observed continuous trait distribution.

For each observed orthogroup statistic, the calibrated p-value is based on the number of matched null statistics more extreme than the observed statistic, with tied statistics given half weight and a small finite-sample correction:

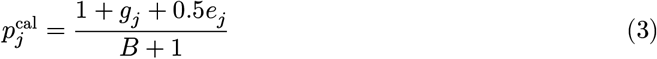

For two-sided tests, more extreme statistics are evaluated using |*Z*|. Here, *B* is the number of matched null statistics, *g*_*j*_ is the number more extreme than the observed statistic, and *e*_*j*_ is the number exactly tied with it. The calibrated p-values are then used for Benjamini-Hochberg FDR correction and final association calls.

Calibration is performed separately for each test mode. Presence/absence tests are calibrated within prevalence-matched representative bins, so each observed orthogroup is compared against null orthogroups with similar occupancy across species. Copy-number tests use a mode-specific global null pool or matched mean-count/sparsity bins, depending on the production setting. OrthoGLMM reports calibration null-pool size, KS distance from Uniform(0, 1), tail lambda, empirical p-value floor behavior, and mode-specific QQ summaries.

## 5 Implementation

OrthoGLMM is implemented as a reproducible command-line workflow. Each production run records the run configuration, command log package version, input hashes, output hashes, runtime report, solver metadata, null-task manifests, and calibration summaries. The main outputs are the calibrated genome-wide table, top-candidate table, calibrator JSON, and mode-level QC summaries. Supplemental outputs include the full calibrated orthogroup results, meta-family results when used, calibration settings and null depths by mode or bin, and optional node-localization or sequence-triage outputs.

Optional secondary analyses can group related orthogroups into protein meta-families. Representative proteins are selected from each orthogroup, compared with an all-vs-all protein similarity search, converted into an orthogroup-level graph, and clustered with an algorithm such as MCL. The resulting meta-family features are scanned and calibrated with the same OrthoGLMM framework.

Optional node-localization and sequence-triage tools help interpret significant or high-priority orthogroups. Node-localization tests whether an orthogroup content signal is concentrated in an internal clade of the orthogroup gene tree. Sequence triage computes simple relative-rate summaries for near-single-copy, alignment-suitable orthogroups and routes candidates to dedicated sequence-evolution analyses. Claims about amino-acid convergence, codon-level selection, branch-site effects, or causal molecular convergence require downstream tools designed for those questions.

## 6 Results

### 6.1 Dataset and gene prediction summary

We benchmarked OrthoGLMM across three cohorts and three traits: diazotrophy in bacteria, nodulation in plants, and marine lifestyle in mammals. These benchmarks have well-characterized biological expectations and test whether OrthoGLMM can recover expected signals from *de novo* gene annotation.

**Table 1.** Dataset, orthogroup, prediction summary, and mean BUSCO completeness of predicted gene annotation for the three benchmark analyses.

| Cohort | Trait | Taxa | Positive | Proteins | Orthogroups | BUSCO |
| --- | --- | --- | --- | --- | --- | --- |
| Bacteria | Diazotrophy | 400 | 77 | 1,410,711 | 67,653 | 94.6 |
| Plants | Nodulation | 89 | 60 | 3,156,926 | 87,723 | 59.2 |
| Mammals | Marine lifestyle | 106 | 36 | 3,296,851 | 55,925 | 92.8 |

Full ranked top-hit tables for copy-number and presence/absence analyses are provided in Supplementary Tables S1-S6. Orthogroups are discussed in raw Wald-statistic order; calibrated q-values are computed from mode- and bin-specific empirical nulls and can differ from the raw rank order.

### 6.2 Diazotrophy in bacteria

Gene annotation produced an average of 3,874 genes per genome (range 597-10,331; 1,549,771 total predicted proteins), of which 1,410,711 were placed into 67,653 orthogroups. Of the 400 bacterial species analysed, 77 were positive for diazotrophy (19.25%). Only 18 orthogroups were present in all 400 species, reflecting high gene-content diversity across the dataset. The complete species list is provided in Supplementary Table S7.

Diazotrophy is the most direct positive-control benchmark because the expected signal is encoded in gene content. The strongest associated orthogroups recovered components of the canonical molybdenum nitrogenase system and FeMo-cofactor assembly pathway^21,73–76^. These included the structural nitrogenase genes nifH, nifD, and nifK, together with cofactor and accessory functions including nifE, nifB, nifV, nifQ, and nifS-like proteins.

Additional expected nitrogen-fixation-associated genes were recovered at lower ranks, including nifS, nifU, nifX, hyp/hup hydrogenase maturation genes, molybdenum/cofactor-related genes such as modB and mopI, and fixation/redox-associated genes such as fixA and cowN. This ranking is expected because many accessory, maturation, transport, hydrogenase, and redox genes are biologically relevant to diazotrophy and also occur in broader metabolic contexts than core nitrogenase structural and FeMo-cofactor genes. Rank is therefore a genome-wide association ordering; pathway importance should be interpreted from biological context and follow-up evidence.

### 6.3 Nodulation in plants

Gene annotation produced an average of 35,471 genes per genome (range 17,518-94,264; 3,156,926 total predicted proteins), which were placed into 87,723 orthogroups. Of the 89 plant species analysed, 60 were positive for nodulation (67.42%). A total of 5,371 orthogroups were present in all species, reflecting a substantial conserved plant gene complement across the dataset. The complete species list is provided in Supplementary Table S8.

In the nodulation benchmark, nitrogenase is encoded by the bacterial symbiont. The expected plant signal is therefore host-side nodulation biology, including rhizobial recognition and infection, nodule development, cell-wall remodeling, transport, carbon metabolism, and oxygen/redox physiology^77,78^. The copy-number analysis recovered several high-ranking functions in these well-studied categories, including annexin, sugar transport, malate transport, pectinesterase/ pectinesterase inhibitor, fasciclin-like arabinogalactan protein, and hemoglobin-family genes^79–83^. Non-symbiotic hemoglobin and two-on-two hemoglobin-3 ranked 4 and 8, respectively, consistent with the central role of hemoglobin-family proteins in nodule oxygen and redox physiology.

The presence/absence analysis also recovered direct nodulation-associated functions. Early nodulin-12A was among the strongest positive associations, and multiple fasciclin-like arabinogalactan protein orthogroups were enriched in nodulating species. ENOD12 is a well-studied early nodulation marker, while arabinogalactan/extensin-type glycoproteins are associated with infection-thread and symbiotic-interface cell-wall biology^82,84^. The plant benchmark recovered a coherent host-side nodulation signal across copy-number and presence/absence modes, with strongest support for infection-associated cell-wall biology, transport and carbon supply, and oxygen/redox physiology.

### 6.4 Marine lifestyle in mammals

Gene annotation produced an average of 31,630 genes per genome (range 25,642-45,311; 3,352,729 total predicted proteins), of which 3,296,851 were placed into 55,925 orthogroups. Of the 106 mammalian species analysed, 36 were positive for marine lifestyle (33.96%). A total of 5,050 orthogroups were present in all species. The complete species list is provided in Supplementary Table S9.

The strongest marine-associated orthogroups recovered the expected contraction of chemosensory gene families in secondarily aquatic mammals^30^. These included multiple olfactory receptor orthogroups, bitter taste receptor orthogroups including TAS2R4 and TAS2R1, and the trace amine-associated receptor TAAR5. Most high-ranked receptor orthogroups had negative effects, indicating reduced copy number or depletion in marine species, consistent with reduced olfactory receptor repertoires and altered taste and chemosensory biology in aquatic lineages.

Additional high-ranking signals included skin, hair, epithelial barrier, and secretory functions, such as PIP, KRT10, KRT1, KRT35, KRT72, other keratin-associated proteins, and SERPINB7, as well as lipid, mitochondrial, or muscle-energy genes such as LDHB, ATP synthase, and tricarboxylate transport orthogroups. These categories are plausible marine-lifestyle candidates, although many classic marine mammal adaptations involve sequence evolution, regulation, morphology, or physiology beyond gene gain/loss. The mammal scan therefore recovers a strong gene-content signal for chemosensory receptor depletion, with additional candidate signals in epithelial, metabolic, and sensory categories.

## 7 Discussion

OrthoGLMM provides a phylogenetically informed framework for genome-wide association between species-level traits and gene-content features. It scales association testing across orthogroup presence/absence and copy-number matrices, while using empirical solver-rerun calibration to reduce reliance on nominal Wald statistics. This is useful for comparative datasets where orthogroups vary widely in prevalence, sparsity, copy-number distribution, and phylogenetic clustering.

The benchmark analyses show the range of expected use cases. Diazotrophy in bacteria provides a direct positive control because the focal biology is encoded in gene content; OrthoGLMM recovered nitrogenase structural genes and FeMo-cofactor biosynthesis genes among the strongest associations. For plant nodulation, the recovered signals reflect host-side oxygen/redox physiology, cell-wall modification, transport, and infection-associated functions because nitrogenase is encoded by the bacterial symbiont. Marine lifestyle in mammals showed the clearest signal in chemosensory receptor depletion, with additional candidate signals in epithelial, lipid/metabolic, and sensory categories.

Calibration is central to interpreting these scans. Orthogroup data include rare, nearly fixed, clade-restricted, and phylogenetically correlated features, making raw Wald p-values unreliable as final evidence. OrthoGLMM therefore uses raw statistics for ranking and diagnostics, then estimates calibrated p-values and FDR from matched solver-rerun null distributions.

The main limitations come from upstream data and biological interpretation. Assembly, annotation, orthogroup inference, trait coding, covariate choice, and tree uncertainty can all affect results. Broad orthogroups can hide clade-specific signals, while narrow orthogroups can split one biological family across multiple tests. HOG, node-localization, meta-family, and sequence-triage layers can help prioritize candidates; functional mechanism still requires follow-up with expression data, domain annotation, experimental validation, or sequence-evolution models.

OrthoGLMM is best used as a calibrated comparative screen for candidate discovery. It lets researchers search broad genome datasets, prioritize candidate gene families, and generate focused hypotheses for experimental, functional, or sequence-level validation. Future development should prioritize domain-level features, stronger HOG and meta-family summaries, multistate and continuous traits, and clearer visualization of calibration behavior and candidate-gene evidence.

## 8 Conclusion

OrthoGLMM performs phylogenetically informed genome-content association scans across orthogroup presence/absence and copy-number features. In three benchmark datasets, it recovered expected signals for bacterial diazotrophy, plant nodulation, and chemosensory receptor depletion in marine mammals. Solver-rerun empirical calibration improves interpretation under phylogenetic structure and sparsity, while optional follow-up layers help prioritize candidates for downstream functional or sequence-evolution analyses.

## Supporting information

Supplemental Dta

## 9 Data and software availability

Source code, documentation, example data, and reproducibility scripts will be made available at http://github.com/jguhlin/OrthoGLMM. Genome accessions, trait coding, full calibrated outputs, and supplementary tables are provided as Supplementary Data.

