## Supplemental Dta for "OrthoGLMM: Phylogenetic Association Testing for Gene Content and Trait Evolution"

### Supplementary Information

Top 50 ranked top-hit tables for copy-number and presence/absence analyses are provided in Supplementary Tables S1-S6.

#### Supplementary table index

- **S1.** Bacterial diazotrophy, copy-number: top 50 ranked copy-number association hits for diazotrophy across bacterial orthogroups.
- **S2.** Bacterial diazotrophy, presence/absence: top 50 ranked presence/absence association hits for diazotrophy across bacterial orthogroups.
- **S3.** Plant nodulation, copy-number: top 50 ranked copy-number association hits for nodulation across plant orthogroups.
- **S4.** Plant nodulation, presence/absence: top 50 ranked presence/absence association hits for nodulation across plant orthogroups.
- **S5.** Mammalian marine lifestyle, copy-number: top 50 ranked copy-number association hits for marine lifestyle across mammalian orthogroups.
- **S6.** Mammalian marine lifestyle, presence/absence: top 50 ranked presence/absence association hits for marine lifestyle across mammalian orthogroups.
- **S7.** Bacterial species list: complete list of bacterial species analysed, including diazotrophy labels and associated metadata.
- **S8.** Plant species list: complete list of plant species analysed, including nodulation labels and associated metadata.
- **S9.** Mammalian species list: complete list of mammalian species analysed, including marine lifestyle labels and associated metadata.

#### Bacterial diazotrophy

Gene annotation produced an average of 3,874 genes per genome (range 597-10,331; 1,549,771 total predicted proteins), of which 1,410,711 were placed into 67,653 orthogroups. Of the 400 bacterial species analysed, 77 were positive for diazotrophy (19.25%). Only 18 orthogroups were present in all 400 species, reflecting high gene-content diversity across the dataset. The complete species list is provided in Supplementary Table S7.

#### Plant nodulation

The plant analysis comprised 89 species, of which 60 were positive for nodulation (67.42%). A total of 5,371 orthogroups were present in all species, reflecting a substantial conserved plant gene complement across the dataset. The complete species list is provided in Supplementary Table S8.

#### Marine lifestyle in mammals

Gene annotation produced an average of 47,945 genes per genome (range 25,642-138,074; 5,082,182 total predicted proteins), all of which were placed into 81,659 orthogroups. Of the 106 mammalian species analysed, 36 were positive for marine lifestyle (33.96%). A total of 7,626 orthogroups were present in all species. The complete species list is provided in Supplementary Table S9.

#### Supplementary Table S1

Top 50 ranked copy-number association hits for diazotrophy across 400 bacterial species. Rows correspond to orthogroups and include rank, orthogroup identifier, UniProt hit, putative function, effect direction, effect estimate, test statistic, raw p value, calibrated p value, false-discovery rate, prevalence, and mean copy number.

| Rank | Orthogroup | UniProt | Putative_function | Direction | Beta | z | raw_p | cal_p | FDR | Prev | Mean_CN |
|---|---|---|---|---|---|---|---|---|---|---|---|
| 1 | OG0005550 | P20621 (NIFK_BRADU) | Nitrogenase molybdenum-iron protein beta chain | enriched | 1.38 | 11.2 | 7.16e-29 | 1.43e-05 | 0.0279 | 0.147 | 0.155 |
| 2 | OG0007589 | Q44491 (Y4254_TRIV2) | UPF0437 protein Ava_4254 | enriched | 1.37 | 11.1 | 1.16e-28 | 1.43e-05 | 0.0279 | 0.115 | 0.117 |
| 3 | OG0005316 | Q47921 (NIFH2_MASLA) | Nitrogenase iron protein 2 | enriched | 1.34 | 10.8 | 2.77e-27 | 1.43e-05 | 0.0279 | 0.145 | 0.16 |
| 4 | OG0001951 | P14889 (NIFZ_AZOVI) | Protein NifZ | enriched | 1.33 | 10.8 | 3.85e-27 | 1.43e-05 | 0.0279 | 0.138 | 0.34 |
| 5 | OG0006745 | P14888 (NIFW_AZOVI) | Nitrogenase-stabilizing/protective protein NifW | enriched | 1.33 | 10.8 | 5.91e-27 | 1.43e-05 | 0.0279 | 0.13 | 0.133 |
| 6 | OG0005469 | P58567 (FER3_NOSS1) | Ferredoxin-3 | enriched | 1.32 | 10.7 | 1.06e-26 | 1.43e-05 | 0.0279 | 0.145 | 0.158 |
| 7 | OG0005057 | P14886 (NIFY_AZOVI) | Protein NifY | enriched | 1.33 | 10.7 | 1.55e-26 | 1.43e-05 | 0.0279 | 0.145 | 0.168 |
| 8 | OG0006328 | P55676 (Y4VQ_SINFN) | UPF0460 protein y4vQ | enriched | 1.3 | 10.6 | 3.02e-26 | 1.43e-05 | 0.0279 | 0.133 | 0.14 |
| 9 | OG0002798 | Q44144 (NIFE_NOSS1) | Nitrogenase iron-molybdenum cofactor biosynthesis protein NifE | enriched | 1.25 | 9.96 | 2.19e-23 | 1.43e-05 | 0.0279 | 0.24 | 0.263 |
| 10 | OG0003265 | P00467 (NIFD_CLOPA) | Nitrogenase molybdenum-iron protein alpha chain | enriched | 1.24 | 9.81 | 1.03e-22 | 1.43e-05 | 0.0279 | 0.23 | 0.235 |
| 11 | OG0003214 | P46044 (NIFB_FRAAL) | FeMo cofactor biosynthesis protein NifB | enriched | 1.21 | 9.71 | 2.63e-22 | 1.43e-05 | 0.0279 | 0.22 | 0.237 |
| 12 | OG0008681 | C0MAL8 (THSA_STRE4) | NAD(+) hydrolase ThsA | enriched | 1.14 | 9.42 | 4.61e-21 | 1.43e-05 | 0.0279 | 0.102 | 0.102 |
| 13 | OG0007577 | Q53213 (NIFQ_SINFN) | Protein NifQ homolog | enriched | 1.13 | 9.25 | 2.19e-20 | 1.43e-05 | 0.0279 | 0.115 | 0.117 |
| 14 | OG0006100 | P23177 (YNIW_AZOCH) | Uncharacterized 19.8 kDa protein in nifW 5'region | enriched | 1.08 | 9.06 | 1.35e-19 | 1.43e-05 | 0.0279 | 0.095 | 0.145 |
| 15 | OG0005150 | Q01181 (NIFV_CERSP) | Homocitrate synthase | enriched | 1.11 | 8.86 | 7.91e-19 | 1.43e-05 | 0.0279 | 0.165 | 0.165 |
| 16 | OG0000651 | P24496 (FER_SACER) | Ferredoxin | enriched | 1.16 | 8.82 | 1.16e-18 | 1.43e-05 | 0.0279 | 0.395 | 0.605 |
| 17 | OG0002981 | P11067 (NIFB_AZOVI) | FeMo cofactor biosynthesis protein NifB | enriched | 1.12 | 8.81 | 1.22e-18 | 1.43e-05 | 0.0279 | 0.235 | 0.25 |
| 18 | OG0011172 | G3IZ31 (G3IZ31_METTV) | DUF6129 domain-containing protein | enriched | 1.04 | 8.69 | 3.68e-18 | 1.43e-05 | 0.0279 | 0.0775 | 0.0775 |
| 19 | OG0000405 | P29847 (CYSE_SALTY) | Serine acetyltransferase | enriched | 1.16 | 8.42 | 3.91e-17 | 1.43e-05 | 0.0279 | 0.64 | 0.762 |
| 20 | OG0000036 | A4WDA9 (ISCA_ENT38) | Iron-binding protein IscA | enriched | 1.19 | 8.26 | 1.44e-16 | 1.43e-05 | 0.0279 | 0.665 | 1.27 |
| 21 | OG0000382 | P77409 (YDHU_ECOLI) | Putative cytochrome YdhU | enriched | 1.01 | 7.8 | 6.05e-15 | 1.43e-05 | 0.0279 | 0.4 | 0.78 |
| 22 | OG0010115 | P17435 (YNF1_RHOCA) | Uncharacterized 27.7 kDa protein in nifB 3'region | enriched | 0.919 | 7.78 | 7.16e-15 | 1.43e-05 | 0.0279 | 0.0725 | 0.0875 |
| 23 | OG0012208 | A1K639 (COWN_AZOSB) | N(2)-fixation sustaining protein CowN | enriched | 0.903 | 7.54 | 4.88e-14 | 1.43e-05 | 0.0279 | 0.07 | 0.07 |
| 24 | OG0002961 | A2SFV1 (RBL1A_METPP) | Ribulose bisphosphate carboxylase large chain 1 | enriched | 0.918 | 7.43 | 1.12e-13 | 1.43e-05 | 0.0279 | 0.195 | 0.253 |
| 25 | OG0011825 | Q9CD83 (PHBS_MYCLE) | Chorismate pyruvate-lyase | enriched | 0.976 | 7.4 | 1.32e-13 | 1.43e-05 | 0.0279 | 0.065 | 0.0725 |
| 26 | OG0006872 | P44679 (YBGC_HAEIN) | Acyl-CoA thioesterase YbgC | enriched | 0.895 | 7.39 | 1.43e-13 | 1.43e-05 | 0.0279 | 0.085 | 0.13 |
| 27 | OG0006803 | P10331 (FIXC_BRADU) | Protein FixC | enriched | 0.93 | 7.39 | 1.48e-13 | 1.43e-05 | 0.0279 | 0.107 | 0.13 |
| 28 | OG0002020 | P31878 (HUPK_AZOVI) | Hydrogenase expression/formation protein HupK | enriched | 0.912 | 7.3 | 2.95e-13 | 1.43e-05 | 0.0279 | 0.242 | 0.333 |
| 29 | OG0001134 | P82802 (FER2_AZOVD) | Ferredoxin, 2Fe-2s | enriched | 0.937 | 7.27 | 3.6e-13 | 1.43e-05 | 0.0279 | 0.335 | 0.46 |
| 30 | OG0003713 | P37733 (MODE_AZOVI) | Molybdenum transport protein ModE | enriched | 0.92 | 7.23 | 4.97e-13 | 1.43e-05 | 0.0279 | 0.152 | 0.215 |
| 31 | OG0001953 | Q9F1R2 (CMPR_SYNE7) | HTH-type transcriptional activator CmpR | enriched | 0.884 | 7 | 2.52e-12 | 1.43e-05 | 0.0279 | 0.255 | 0.338 |
| 32 | OG0012967 | P14887 (NIFX_AZOVI) | Protein NifX | enriched | 0.841 | 6.97 | 3.13e-12 | 1.43e-05 | 0.0279 | 0.0575 | 0.0625 |
| 33 | OG0004786 | P05341 (NIFS_AZOVI) | Cysteine desulfurase NifS | enriched | 0.865 | 6.91 | 4.73e-12 | 1.43e-05 | 0.0279 | 0.17 | 0.175 |
| 34 | OG0005462 | P31902 (HYPB_CUPNH) | Hydrogenase maturation factor HypB | enriched | 0.856 | 6.89 | 5.64e-12 | 1.43e-05 | 0.0279 | 0.15 | 0.158 |
| 35 | OG0009373 | P14299 (DRAT_RHORU) | NAD(+)--dinitrogen-reductase ADP-D-ribosyltransferase | enriched | 0.825 | 6.84 | 7.69e-12 | 1.43e-05 | 0.0279 | 0.0875 | 0.095 |
| 36 | OG0005605 | P0AC11 (DHP2_ECOLX) | Dihydropteroate synthase type-2 | enriched | 0.952 | 6.77 | 1.33e-11 | 1.43e-05 | 0.0279 | 0.11 | 0.155 |
| 37 | OG0008043 | P13065 (PHSL_DESBA) | Periplasmic [NiFeSe] hydrogenase large subunit | enriched | 0.814 | 6.67 | 2.62e-11 | 1.43e-05 | 0.0279 | 0.1 | 0.11 |
| 38 | OG0001724 | A3PJL9 (LIPA_CERS1) | Lipoyl synthase | enriched | 1 | 6.66 | 2.7e-11 | 1.43e-05 | 0.0279 | 0.352 | 0.365 |
| 39 | OG0011215 | A0A1N7SXT8 (A0A1N7SXT8_9BURK) | Uncharacterized protein | enriched | 0.779 | 6.65 | 2.9e-11 | 1.43e-05 | 0.0279 | 0.0625 | 0.0775 |
| 40 | OG0000845 | P0AAM3 (HYPC_ECOLI) | Hydrogenase maturation factor HypC | enriched | 0.825 | 6.65 | 2.99e-11 | 1.43e-05 | 0.0279 | 0.41 | 0.54 |
| 41 | OG0001553 | O86235 (Y122B_HAEIN) | Uncharacterized protein HI_1225.1 | enriched | 0.929 | 6.58 | 4.62e-11 | 1.43e-05 | 0.0279 | 0.383 | 0.388 |
| 42 | OG0005853 | F4CQ78 (RBS_PSEUX) | Ribulose bisphosphate carboxylase small subunit | enriched | 0.802 | 6.51 | 7.35e-11 | 1.43e-05 | 0.0279 | 0.122 | 0.15 |
| 43 | OG0001337 | P0C7J3 (XANB_XANCP) | Xanthan biosynthesis protein XanB | enriched | 0.87 | 6.5 | 8.17e-11 | 1.43e-05 | 0.0279 | 0.343 | 0.422 |
| 44 | OG0009482 | P14300 (DRAG_RHORU) | ADP-ribosyl-[dinitrogen reductase] glycohydrolase | enriched | 0.797 | 6.45 | 1.09e-10 | 1.43e-05 | 0.0279 | 0.09 | 0.0925 |
| 45 | OG0001034 | P84253 (MOSB_AZOVD) | Molybdenum storage protein subunit beta | enriched | 0.894 | 6.44 | 1.2e-10 | 1.43e-05 | 0.0279 | 0.43 | 0.485 |
| 46 | OG0004834 | P37097 (FER5_RHOCA) | Ferredoxin-5 | enriched | 0.766 | 6.4 | 1.55e-10 | 1.43e-05 | 0.0279 | 0.125 | 0.175 |
| 47 | OG0006678 | P04952 (MOP1_CLOPA) | Molybdenum-pterin-binding protein 1 | enriched | 0.789 | 6.4 | 1.58e-10 | 1.43e-05 | 0.0279 | 0.11 | 0.133 |
| 48 | OG0001520 | Q59297 (CYB6_CHLTI) | Cytochrome bc complex cytochrome b subunit | enriched | 0.907 | 6.36 | 2.01e-10 | 1.43e-05 | 0.0279 | 0.385 | 0.393 |
| 49 | OG0022504 | A0ABR8J5L7 (A0ABR8J5L7_9NOST) | Uncharacterized protein | enriched | 0.775 | 6.36 | 2.07e-10 | 1.43e-05 | 0.0279 | 0.0275 | 0.0275 |
| 50 | OG0000692 | O52058 (ACCC_ALLVD) | Biotin carboxylase | enriched | 0.948 | 6.35 | 2.17e-10 | 1.43e-05 | 0.0279 | 0.573 | 0.593 |

#### Supplementary Table S2

Top 50 ranked presence/absence association hits for diazotrophy across 400 bacterial species. Rows correspond to orthogroups and include rank, orthogroup identifier, UniProt hit, putative function, effect direction, effect estimate, test statistic, raw p value, calibrated p value, false-discovery rate, prevalence, and mean copy number.

| Rank | Orthogroup | UniProt | Putative_function | Direction | Beta | z | raw_p | cal_p | FDR | Prev | Mean_CN |
|---|---|---|---|---|---|---|---|---|---|---|---|
| 1 | OG0005057 | P14886 (NIFY_AZOVI) | Protein NifY | enriched | 1.41 | 11.4 | 6.75e-30 | 0.0001 | 0.0279 | 0.145 | 0.145 |
| 2 | OG0005550 | P20621 (NIFK_BRADU) | Nitrogenase molybdenum-iron protein beta chain | enriched | 1.39 | 11.2 | 3.01e-29 | 0.0001 | 0.0279 | 0.147 | 0.147 |
| 3 | OG0007589 | Q44491 (Y4254_TRIV2) | UPF0437 protein Ava_4254 | enriched | 1.36 | 11.1 | 1.13e-28 | 0.0001 | 0.0279 | 0.115 | 0.115 |
| 4 | OG0005469 | P58567 (FER3_NOSS1) | Ferredoxin-3 | enriched | 1.36 | 11 | 3.47e-28 | 0.0001 | 0.0279 | 0.145 | 0.145 |
| 5 | OG0006328 | P55676 (Y4VQ_SINFN) | UPF0460 protein y4vQ | enriched | 1.36 | 11 | 4.6e-28 | 0.0001 | 0.0279 | 0.133 | 0.133 |
| 6 | OG0005316 | Q47921 (NIFH2_MASLA) | Nitrogenase iron protein 2 | enriched | 1.35 | 11 | 5.91e-28 | 0.0001 | 0.0279 | 0.145 | 0.145 |
| 7 | OG0006745 | P14888 (NIFW_AZOVI) | Nitrogenase-stabilizing/protective protein NifW | enriched | 1.32 | 10.7 | 7.73e-27 | 0.0001 | 0.0279 | 0.13 | 0.13 |
| 8 | OG0001951 | P14889 (NIFZ_AZOVI) | Protein NifZ | enriched | 1.32 | 10.7 | 1.47e-26 | 0.0001 | 0.0279 | 0.138 | 0.138 |
| 9 | OG0003265 | P00467 (NIFD_CLOPA) | Nitrogenase molybdenum-iron protein alpha chain | enriched | 1.25 | 9.93 | 2.96e-23 | 0.0001 | 0.0279 | 0.23 | 0.23 |
| 10 | OG0006100 | P23177 (YNIW_AZOCH) | Uncharacterized 19.8 kDa protein in nifW 5'region | enriched | 1.2 | 9.9 | 4.33e-23 | 0.0001 | 0.0279 | 0.095 | 0.095 |
| 11 | OG0002798 | Q44144 (NIFE_NOSS1) | Nitrogenase iron-molybdenum cofactor biosynthesis protein NifE | enriched | 1.24 | 9.85 | 6.89e-23 | 0.0001 | 0.0279 | 0.24 | 0.24 |
| 12 | OG0003214 | P46044 (NIFB_FRAAL) | FeMo cofactor biosynthesis protein NifB | enriched | 1.22 | 9.73 | 2.27e-22 | 0.0001 | 0.0279 | 0.22 | 0.22 |
| 13 | OG0002981 | P11067 (NIFB_AZOVI) | FeMo cofactor biosynthesis protein NifB | enriched | 1.19 | 9.48 | 2.58e-21 | 0.0001 | 0.0279 | 0.235 | 0.235 |
| 14 | OG0008681 | C0MAL8 (THSA_STRE4) | NAD(+) hydrolase ThsA | enriched | 1.14 | 9.42 | 4.61e-21 | 0.0001 | 0.0279 | 0.102 | 0.102 |
| 15 | OG0007577 | Q53213 (NIFQ_SINFN) | Protein NifQ homolog | enriched | 1.14 | 9.31 | 1.32e-20 | 0.0001 | 0.0279 | 0.115 | 0.115 |
| 16 | OG0005150 | Q01181 (NIFV_CERSP) | Homocitrate synthase | enriched | 1.11 | 8.86 | 7.91e-19 | 0.0001 | 0.0279 | 0.165 | 0.165 |
| 17 | OG0011172 | G3IZ31 (G3IZ31_METTV) | DUF6129 domain-containing protein | enriched | 1.04 | 8.69 | 3.68e-18 | 0.0001 | 0.0279 | 0.0775 | 0.0775 |
| 18 | OG0010115 | P17435 (YNF1_RHOCA) | Uncharacterized 27.7 kDa protein in nifB 3'region | enriched | 0.957 | 8.04 | 9.26e-16 | 0.0001 | 0.0279 | 0.0725 | 0.0725 |
| 19 | OG0006803 | P10331 (FIXC_BRADU) | Protein FixC | enriched | 0.96 | 7.7 | 1.33e-14 | 0.0001 | 0.0279 | 0.107 | 0.107 |
| 20 | OG0011825 | Q9CD83 (PHBS_MYCLE) | Chorismate pyruvate-lyase | enriched | 0.983 | 7.62 | 2.55e-14 | 0.0001 | 0.0279 | 0.065 | 0.065 |
| 21 | OG0012208 | A1K639 (COWN_AZOSB) | N(2)-fixation sustaining protein CowN | enriched | 0.903 | 7.54 | 4.88e-14 | 0.0001 | 0.0279 | 0.07 | 0.07 |
| 22 | OG0006872 | P44679 (YBGC_HAEIN) | Acyl-CoA thioesterase YbgC | enriched | 0.921 | 7.53 | 5.01e-14 | 0.0001 | 0.0279 | 0.085 | 0.085 |
| 23 | OG0011215 | A0A1N7SXT8 (A0A1N7SXT8_9BURK) | Uncharacterized protein | enriched | 0.868 | 7.28 | 3.44e-13 | 0.0001 | 0.0279 | 0.0625 | 0.0625 |
| 24 | OG0002961 | A2SFV1 (RBL1A_METPP) | Ribulose bisphosphate carboxylase large chain 1 | enriched | 0.91 | 7.18 | 6.81e-13 | 0.0001 | 0.0279 | 0.195 | 0.195 |
| 25 | OG0001953 | Q9F1R2 (CMPR_SYNE7) | HTH-type transcriptional activator CmpR | enriched | 0.919 | 7.15 | 8.89e-13 | 0.0001 | 0.0279 | 0.255 | 0.255 |
| 26 | OG0012967 | P14887 (NIFX_AZOVI) | Protein NifX | enriched | 0.868 | 7.13 | 1.02e-12 | 0.0001 | 0.0279 | 0.0575 | 0.0575 |
| 27 | OG0001017 | A3N2T3 (NGT_ACTP2) | UDP-glucose:protein N-beta-glucosyltransferase | enriched | 0.913 | 7.12 | 1.07e-12 | 0.0001 | 0.0279 | 0.225 | 0.225 |
| 28 | OG0005462 | P31902 (HYPB_CUPNH) | Hydrogenase maturation factor HypB | enriched | 0.885 | 7.06 | 1.7e-12 | 0.0001 | 0.0279 | 0.15 | 0.15 |
| 29 | OG0001724 | A3PJL9 (LIPA_CERS1) | Lipoyl synthase | enriched | 1.07 | 6.96 | 3.29e-12 | 0.0001 | 0.0279 | 0.352 | 0.352 |
| 30 | OG0002456 | P0AG78 (SUBI_ECOLI) | Sulfate-binding protein | enriched | 0.917 | 6.89 | 5.76e-12 | 0.0001 | 0.0279 | 0.258 | 0.258 |
| 31 | OG0009373 | P14299 (DRAT_RHORU) | NAD(+)--dinitrogen-reductase ADP-D-ribosyltransferase | enriched | 0.832 | 6.82 | 8.84e-12 | 0.0001 | 0.0279 | 0.0875 | 0.0875 |
| 32 | OG0001520 | Q59297 (CYB6_CHLTI) | Cytochrome bc complex cytochrome b subunit | enriched | 0.972 | 6.79 | 1.13e-11 | 0.0001 | 0.0279 | 0.385 | 0.385 |
| 33 | OG0009482 | P14300 (DRAG_RHORU) | ADP-ribosyl-[dinitrogen reductase] glycohydrolase | enriched | 0.827 | 6.75 | 1.53e-11 | 0.0001 | 0.0279 | 0.09 | 0.09 |
| 34 | OG0008043 | P13065 (PHSL_DESBA) | Periplasmic [NiFeSe] hydrogenase large subunit | enriched | 0.82 | 6.74 | 1.55e-11 | 0.0001 | 0.0279 | 0.1 | 0.1 |
| 35 | OG0001672 | Q21DY7 (UBIA_SACD2) | 4-hydroxybenzoate octaprenyltransferase | enriched | 1.05 | 6.74 | 1.6e-11 | 0.0001 | 0.0279 | 0.362 | 0.362 |
| 36 | OG0004786 | P05341 (NIFS_AZOVI) | Cysteine desulfurase NifS | enriched | 0.841 | 6.72 | 1.82e-11 | 0.0001 | 0.0279 | 0.17 | 0.17 |
| 37 | OG0005853 | F4CQ78 (RBS_PSEUX) | Ribulose bisphosphate carboxylase small subunit | enriched | 0.831 | 6.67 | 2.55e-11 | 0.0001 | 0.0279 | 0.122 | 0.122 |
| 38 | OG0001553 | O86235 (Y122B_HAEIN) | Uncharacterized protein HI_1225.1 | enriched | 0.938 | 6.61 | 3.88e-11 | 0.0001 | 0.0279 | 0.383 | 0.383 |
| 39 | OG0001616 | O87689 (CBIH_PRIMG) | Cobalt-factor III methyltransferase | enriched | 0.875 | 6.56 | 5.55e-11 | 0.0001 | 0.0279 | 0.37 | 0.37 |
| 40 | OG0002020 | P31878 (HUPK_AZOVI) | Hydrogenase expression/formation protein HupK | enriched | 0.831 | 6.55 | 5.75e-11 | 0.0001 | 0.0279 | 0.242 | 0.242 |
| 41 | OG0001337 | P0C7J3 (XANB_XANCP) | Xanthan biosynthesis protein XanB | enriched | 0.906 | 6.53 | 6.58e-11 | 0.0001 | 0.0279 | 0.343 | 0.343 |
| 42 | OG0001743 | P0AEB0 (CYSW_ECOLI) | Sulfate transport system permease protein CysW | enriched | 0.873 | 6.47 | 9.92e-11 | 0.0001 | 0.0279 | 0.312 | 0.312 |
| 43 | OG0001983 | P63353 (CYSA_BRUME) | Sulfate/thiosulfate import ATP-binding protein CysA | enriched | 0.872 | 6.46 | 1.07e-10 | 0.0001 | 0.0279 | 0.312 | 0.312 |
| 44 | OG0004182 | P29929 (COBN_SINSX) | Aerobic cobaltochelatase subunit CobN | enriched | 0.863 | 6.43 | 1.25e-10 | 0.0001 | 0.0279 | 0.193 | 0.193 |
| 45 | OG0002186 | P41032 (CYST_SALTY) | Sulfate transport system permease protein CysT | enriched | 0.87 | 6.43 | 1.27e-10 | 0.0001 | 0.0279 | 0.28 | 0.28 |
| 46 | OG0003713 | P37733 (MODE_AZOVI) | Molybdenum transport protein ModE | enriched | 0.821 | 6.42 | 1.34e-10 | 0.0001 | 0.0279 | 0.152 | 0.152 |
| 47 | OG0000651 | P24496 (FER_SACER) | Ferredoxin | enriched | 0.89 | 6.41 | 1.44e-10 | 0.0001 | 0.0279 | 0.395 | 0.395 |
| 48 | OG0009043 | O07345 (CHLD_SYNE7) | Magnesium-chelatase subunit ChlD | enriched | 0.785 | 6.4 | 1.56e-10 | 0.0001 | 0.0279 | 0.095 | 0.095 |
| 49 | OG0002370 | P09133 (NIFA_AZOC5) | Nif-specific regulatory protein | enriched | 0.803 | 6.35 | 2.21e-10 | 0.0001 | 0.0279 | 0.22 | 0.22 |
| 50 | OG0008510 | Q8YM64 (CHLN_NOSS1) | Light-independent protochlorophyllide reductase subunit N | enriched | 0.778 | 6.34 | 2.37e-10 | 0.0001 | 0.0279 | 0.102 | 0.102 |

#### Supplementary Table S3

Top 50 ranked copy-number association hits for nodulation across 89 plant species. Rows correspond to orthogroups and include rank, orthogroup identifier, UniProt hit, putative function, effect direction, effect estimate, test statistic, raw p value, calibrated p value, false-discovery rate, prevalence, and mean copy number.

| Rank | Orthogroup | UniProt | Putative_function | Direction | Beta | z | raw_p | cal_p | FDR | Prev | Mean_CN |
|---|---|---|---|---|---|---|---|---|---|---|---|
| 1 | OG0000853 | Q6WWW4 (UPL3_ARATH) | E3 ubiquitin-protein ligase UPL3 | enriched | 1.44 | 5.25 | 1.49e-07 | 3.33e-05 | 0.0264 | 1 | 6.09 |
| 2 | OG0000733 | Q8GY23 (UPL1_ARATH) | E3 ubiquitin-protein ligase UPL1 | enriched | 1.19 | 4.82 | 1.45e-06 | 3.33e-05 | 0.0264 | 1 | 6.57 |
| 3 | OG0009426 | D8WUA4 (SECA2_ARATH) | Protein translocase subunit SECA2, chloroplastic | depleted | -1.31 | -4.74 | 2.09e-06 | 3.33e-05 | 0.0264 | 0.202 | 0.258 |
| 4 | OG0000713 | O24457 (ODPA3_ARATH) | Pyruvate dehydrogenase E1 component subunit alpha-3, chloroplastic | enriched | 1.22 | 4.73 | 2.24e-06 | 3.33e-05 | 0.0264 | 1 | 6.64 |
| 5 | OG0000375 | F4IRU3 (MYO12_ARATH) | Myosin-12 | enriched | 1.28 | 4.56 | 5.21e-06 | 3.33e-05 | 0.0264 | 0.809 | 9.28 |
| 6 | OG0008700 | F4IV99 (CHR5_ARATH) | Protein CHROMATIN REMODELING 5 | depleted | -1.22 | -4.44 | 9.2e-06 | 3.33e-05 | 0.0264 | 0.258 | 0.528 |
| 7 | OG0002071 | Q4F883 (SG101_ARATH) | Senescence-associated carboxylesterase 101 | enriched | 1.17 | 4.43 | 9.42e-06 | 3.33e-05 | 0.0264 | 1 | 3.84 |
| 8 | OG0004648 | Q9T048 (DRL27_ARATH) | Disease resistance protein At4g27190 | enriched | 1.05 | 4.43 | 9.58e-06 | 3.33e-05 | 0.0264 | 1 | 2.07 |
| 9 | OG0010162 | B3LF48 (EHD2_ARATH) | EH domain-containing protein 2 | depleted | -1.08 | -4.36 | 1.32e-05 | 3.33e-05 | 0.0264 | 0.169 | 0.18 |
| 10 | OG0009001 | P92523 (M860_ARATH) | Uncharacterized mitochondrial protein AtMg00860 | depleted | -1.1 | -4.34 | 1.42e-05 | 3.33e-05 | 0.0264 | 0.258 | 0.36 |
| 11 | OG0002275 | F4J7T2 (EAF1B_ARATH) | Chromatin modification-related protein EAF1 B | enriched | 1.12 | 4.28 | 1.9e-05 | 3.33e-05 | 0.0264 | 0.989 | 3.61 |
| 12 | OG0001326 | Q6IMT1 (SAB_ARATH) | Protein SABRE | enriched | 1.03 | 4.24 | 2.19e-05 | 3.33e-05 | 0.0264 | 1 | 4.85 |
| 13 | OG0000222 | A4GGE1 (YCF2_PHAVU) | Protein Ycf2 | enriched | 1.04 | 4.2 | 2.65e-05 | 3.33e-05 | 0.0264 | 1 | 12.4 |
| 14 | OG0004970 | Q94HW2 (POLR1_ARATH) | Retrovirus-related Pol polyprotein from transposon RE1 | enriched | 1.06 | 4.16 | 3.21e-05 | 3.33e-05 | 0.0264 | 0.955 | 1.96 |
| 15 | OG0008760 | Q9LIK7 (ACA13_ARATH) | Putative calcium-transporting ATPase 13, plasma membrane-type | depleted | -0.99 | -4.14 | 3.42e-05 | 3.33e-05 | 0.0264 | 0.281 | 0.483 |
| 16 | OG0001544 | Q76CU2 (PDR1_TOBAC) | Pleiotropic drug resistance protein 1 | enriched | 1.13 | 4.09 | 4.23e-05 | 3.33e-05 | 0.0264 | 0.831 | 4.49 |
| 17 | OG0000728 | F4IG73 (BCHC2_ARATH) | BEACH domain-containing protein C2 | enriched | 1.07 | 4.03 | 5.53e-05 | 3.33e-05 | 0.0264 | 1 | 6.61 |
| 18 | OG0008873 | Q9LHP4 (RGI1_ARATH) | LRR receptor-like serine/threonine-protein kinase RGI1 | depleted | -0.954 | -4.03 | 5.55e-05 | 3.33e-05 | 0.0264 | 0.281 | 0.416 |
| 19 | OG0001614 | F4I9A2 (TNO1_ARATH) | Trans-Golgi network-localized SYP41-interacting protein 1 | enriched | 0.987 | 3.98 | 6.78e-05 | 3.33e-05 | 0.0264 | 1 | 4.39 |
| 20 | OG0001566 | P93295 (M310_ARATH) | Uncharacterized mitochondrial protein AtMg00310 | enriched | 1.04 | 3.97 | 7.16e-05 | 3.33e-05 | 0.0264 | 1 | 4.46 |
| 21 | OG0009153 | Q94HW2 (POLR1_ARATH) | Retrovirus-related Pol polyprotein from transposon RE1 | depleted | -1.01 | -3.96 | 7.64e-05 | 3.33e-05 | 0.0264 | 0.258 | 0.315 |
| 22 | OG0004326 | F4J7T3 (EAF1A_ARATH) | Chromatin modification-related protein EAF1 A | enriched | 0.991 | 3.95 | 7.86e-05 | 3.33e-05 | 0.0264 | 1 | 2.21 |
| 23 | OG0000313 | Q9FR53 (TOR_ARATH) | Serine/threonine-protein kinase TOR | enriched | 0.955 | 3.94 | 8.31e-05 | 3.33e-05 | 0.0264 | 0.989 | 10.3 |
| 24 | OG0001912 | F4I718 (CSI3_ARATH) | Protein CELLULOSE SYNTHASE INTERACTIVE 3 | enriched | 0.977 | 3.93 | 8.51e-05 | 3.33e-05 | 0.0264 | 1 | 4.03 |
| 25 | OG0008006 | Q94HW2 (POLR1_ARATH) | Retrovirus-related Pol polyprotein from transposon RE1 | enriched | 1 | 3.93 | 8.61e-05 | 3.33e-05 | 0.0264 | 0.843 | 1.22 |
| 26 | OG0001151 | Q6IMT1 (SAB_ARATH) | Protein SABRE | enriched | 1.03 | 3.93 | 8.66e-05 | 3.33e-05 | 0.0264 | 1 | 5.22 |
| 27 | OG0004426 | F4HZB2 (BCHA1_ARATH) | Protein SPIRRIG | enriched | 0.968 | 3.92 | 8.76e-05 | 3.33e-05 | 0.0264 | 1 | 2.17 |
| 28 | OG0000488 | Q40253 (L6_LINUS) | Disease resistance protein L6 | enriched | 1.03 | 3.92 | 8.81e-05 | 3.33e-05 | 0.0264 | 0.82 | 8.02 |
| 29 | OG0000710 | P31843 (RRPO_OENBE) | RNA-directed DNA polymerase homolog | enriched | 0.957 | 3.9 | 9.64e-05 | 3.33e-05 | 0.0264 | 1 | 6.66 |
| 30 | OG0000440 | Q9C8Z4 (HEAT1_ARATH) | Uncharacterized protein At3g06530 | enriched | 0.976 | 3.9 | 9.67e-05 | 3.33e-05 | 0.0264 | 1 | 8.39 |
| 31 | OG0004337 | Q8GY23 (UPL1_ARATH) | E3 ubiquitin-protein ligase UPL1 | enriched | 0.987 | 3.89 | 9.83e-05 | 3.33e-05 | 0.0264 | 1 | 2.2 |
| 32 | OG0008715 | P92516 (M750_ARATH) | Uncharacterized mitochondrial protein AtMg00750 | depleted | -1.09 | -3.89 | 0.000101 | 3.33e-05 | 0.0264 | 0.36 | 0.506 |
| 33 | OG0000277 | Q8GY23 (UPL1_ARATH) | E3 ubiquitin-protein ligase UPL1 | enriched | 1.03 | 3.88 | 0.000104 | 3.33e-05 | 0.0264 | 1 | 11 |
| 34 | OG0004094 | P31843 (RRPO_OENBE) | RNA-directed DNA polymerase homolog | enriched | 1.11 | 3.84 | 0.000123 | 3.33e-05 | 0.0264 | 0.618 | 2.31 |
| 35 | OG0001390 | Q9T048 (DRL27_ARATH) | Disease resistance protein At4g27190 | enriched | 0.931 | 3.83 | 0.000129 | 3.33e-05 | 0.0264 | 1 | 4.73 |
| 36 | OG0003854 | Q94KE2 (TIC_ARATH) | Protein TIME FOR COFFEE | enriched | 0.956 | 3.83 | 0.000129 | 3.33e-05 | 0.0264 | 1 | 2.43 |
| 37 | OG0003630 | P31843 (RRPO_OENBE) | RNA-directed DNA polymerase homolog | enriched | 0.932 | 3.82 | 0.000133 | 3.33e-05 | 0.0264 | 0.989 | 2.54 |
| 38 | OG0000112 | F4IHS2 (SYD_ARATH) | Chromatin structure-remodeling complex protein SYD | enriched | 0.946 | 3.8 | 0.000144 | 3.33e-05 | 0.0264 | 1 | 18.5 |
| 39 | OG0002807 | F4JIF4 (NET1B_ARATH) | Protein NETWORKED 1B | enriched | 1.08 | 3.8 | 0.000146 | 3.33e-05 | 0.0264 | 1 | 3.07 |
| 40 | OG0006493 | Q0WL80 (UGGG_ARATH) | UDP-glucose:glycoprotein glucosyltransferase | enriched | 0.962 | 3.77 | 0.000161 | 3.33e-05 | 0.0264 | 0.944 | 1.56 |
| 41 | OG0000509 | Q8GY23 (UPL1_ARATH) | E3 ubiquitin-protein ligase UPL1 | enriched | 0.943 | 3.77 | 0.000165 | 3.33e-05 | 0.0264 | 0.989 | 7.87 |
| 42 | OG0009572 | Q2R2D5 (XA21_ORYSJ) | Receptor kinase-like protein Xa21 | depleted | -0.982 | -3.75 | 0.000179 | 3.33e-05 | 0.0264 | 0.18 | 0.236 |
| 43 | OG0003512 | Q8VY10 (UBC24_ARATH) | Probable ubiquitin-conjugating enzyme E2 24 | enriched | 0.915 | 3.75 | 0.00018 | 3.33e-05 | 0.0264 | 1 | 2.61 |
| 44 | OG0004266 | Q766Z3 (REV3_ARATH) | DNA polymerase zeta catalytic subunit | enriched | 0.962 | 3.74 | 0.000184 | 3.33e-05 | 0.0264 | 1 | 2.24 |
| 45 | OG0004404 | Q9SSD2 (PRP8A_ARATH) | Pre-mRNA-processing-splicing factor 8A | enriched | 0.992 | 3.74 | 0.000185 | 3.33e-05 | 0.0264 | 0.989 | 2.18 |
| 46 | OG0003633 | Q9SSD2 (PRP8A_ARATH) | Pre-mRNA-processing-splicing factor 8A | enriched | 0.95 | 3.71 | 0.00021 | 3.33e-05 | 0.0264 | 1 | 2.54 |
| 47 | OG0001248 | F4KBP5 (CHR4_ARATH) | Protein CHROMATIN REMODELING 4 | enriched | 0.97 | 3.71 | 0.000211 | 3.33e-05 | 0.0264 | 0.989 | 5.02 |
| 48 | OG0004960 | Q8LFJ9 (GLIP7_ARATH) | GDSL esterase/lipase 7 | enriched | 0.906 | 3.7 | 0.000213 | 3.33e-05 | 0.0264 | 1 | 1.97 |
| 49 | OG0000337 | Q9FKS4 (ATR_ARATH) | Serine/threonine-protein kinase ATR | enriched | 0.89 | 3.69 | 0.000224 | 3.33e-05 | 0.0264 | 1 | 9.81 |
| 50 | OG0000454 | Q9FKS4 (ATR_ARATH) | Serine/threonine-protein kinase ATR | enriched | 0.896 | 3.69 | 0.000225 | 3.33e-05 | 0.0264 | 1 | 8.29 |

### Supplementary Table S4

Top 50 ranked presence/absence association hits for nodulation across 89 plant species. Rows correspond to orthogroups and include rank, orthogroup identifier, UniProt hit, putative function, effect direction, effect estimate, test statistic, raw p value, calibrated p value, false-discovery rate, prevalence, and mean copy number.

| Rank | Orthogroup | UniProt | Putative_function | Direction | Beta | z | raw_p | cal_p | FDR | Prev | Mean_CN |
|---|---|---|---|---|---|---|---|---|---|---|---|
| 1 | OG0006111 | F4I096 (MED13_ARATH) | Mediator of RNA polymerase II transcription subunit 13 | depleted | -1.6 | -5.57 | 2.57e-08 | 0.0001 | 0.0264 | 0.494 | 0.494 |
| 2 | OG0004094 | P31843 (RRPO_OENBE) | RNA-directed DNA polymerase homolog | enriched | 1.62 | 5.49 | 4.04e-08 | 0.0001 | 0.0264 | 0.618 | 0.618 |
| 3 | OG0008771 | F4HZB2 (BCHA1_ARATH) | Protein SPIRRIG | depleted | -1.41 | -4.85 | 1.21e-06 | 0.0001 | 0.0264 | 0.348 | 0.348 |
| 4 | OG0005229 | P0C2F6 (RNHX1_ARATH) | Putative ribonuclease H protein At1g65750 | depleted | -1.21 | -4.73 | 2.23e-06 | 0.0001 | 0.0264 | 0.427 | 0.427 |
| 5 | OG0008514 | F4I096 (MED13_ARATH) | Mediator of RNA polymerase II transcription subunit 13 | depleted | -1.26 | -4.6 | 4.24e-06 | 0.0001 | 0.0264 | 0.528 | 0.528 |
| 6 | OG0010162 | B3LF48 (EHD2_ARATH) | EH domain-containing protein 2 | depleted | -1.15 | -4.54 | 5.51e-06 | 0.0001 | 0.0264 | 0.169 | 0.169 |
| 7 | OG0006665 | Q6JN46 (EIX2_SOLLC) | Receptor-like protein EIX2 | depleted | -1.13 | -4.52 | 6.09e-06 | 0.0001 | 0.0264 | 0.157 | 0.157 |
| 8 | OG0009194 | F4IAT2 (THOC2_ARATH) | THO complex subunit 2 | depleted | -1.29 | -4.49 | 7.04e-06 | 0.0001 | 0.0264 | 0.18 | 0.18 |
| 9 | OG0009246 | Q40392 (TMVRN_NICGU) | TMV resistance protein N | depleted | -1.29 | -4.49 | 7.04e-06 | 0.0001 | 0.0264 | 0.18 | 0.18 |
| 10 | OG0009572 | Q2R2D5 (XA21_ORYSJ) | Receptor kinase-like protein Xa21 | depleted | -1.29 | -4.49 | 7.04e-06 | 0.0001 | 0.0264 | 0.18 | 0.18 |
| 11 | OG0008272 | Q9SJQ6 (ROS1_ARATH) | DNA glycosylase/AP lyase ROS1 | depleted | -1.33 | -4.4 | 1.1e-05 | 0.0001 | 0.0264 | 0.528 | 0.528 |
| 12 | OG0008477 | B6EUB3 (PDS5A_ARATH) | Sister chromatid cohesion protein PDS5 homolog A | depleted | -1.19 | -4.39 | 1.16e-05 | 0.0001 | 0.0264 | 0.472 | 0.472 |
| 13 | OG0009505 | Q7PC88 (AB31G_ARATH) | ABC transporter G family member 31 | depleted | -1.13 | -4.34 | 1.46e-05 | 0.0001 | 0.0264 | 0.157 | 0.157 |
| 14 | OG0000216 | Q8GY23 (UPL1_ARATH) | E3 ubiquitin-protein ligase UPL1 | enriched | 1.2 | 4.27 | 1.95e-05 | 0.0001 | 0.0264 | 0.854 | 0.854 |
| 15 | OG0009917 | Q9STE7 (R13L3_ARATH) | Putative disease resistance RPP13-like protein 3 | depleted | -1.12 | -4.22 | 2.42e-05 | 0.0001 | 0.0264 | 0.146 | 0.146 |
| 16 | OG0008936 | F4JTS8 (NOV_ARATH) | Protein NO VEIN | depleted | -1.19 | -4.21 | 2.57e-05 | 0.0001 | 0.0264 | 0.303 | 0.303 |
| 17 | OG0004739 | F4IN58 (PIEZO_ARATH) | Piezo-type mechanosensitive ion channel homolog | depleted | -1.21 | -4.2 | 2.66e-05 | 0.0001 | 0.0264 | 0.146 | 0.146 |
| 18 | OG0008525 | P31843 (RRPO_OENBE) | RNA-directed DNA polymerase homolog | depleted | -1.16 | -4.19 | 2.78e-05 | 0.0001 | 0.0264 | 0.303 | 0.303 |
| 19 | OG0009574 | Q9FWX7 (AB11B_ARATH) | ABC transporter B family member 11 | depleted | -1.18 | -4.19 | 2.8e-05 | 0.0001 | 0.0264 | 0.169 | 0.169 |
| 20 | OG0009652 | Q03194 (PMA4_NICPL) | Plasma membrane ATPase 4 | depleted | -1.18 | -4.19 | 2.8e-05 | 0.0001 | 0.0264 | 0.169 | 0.169 |
| 21 | OG0009441 | Q9LV10 (PTR53_ARATH) | Protein NRT1/ PTR FAMILY 2.11 | depleted | -1.19 | -4.16 | 3.22e-05 | 0.0001 | 0.0264 | 0.146 | 0.146 |
| 22 | OG0008530 | Q60CZ8 (R1A10_SOLDE) | Putative late blight resistance protein homolog R1A-10 | depleted | -1.2 | -4.08 | 4.46e-05 | 0.0001 | 0.0264 | 0.337 | 0.337 |
| 23 | OG0008573 | P92523 (M860_ARATH) | Uncharacterized mitochondrial protein AtMg00860 | depleted | -0.932 | -4.03 | 5.47e-05 | 0.0001 | 0.0264 | 0.18 | 0.18 |
| 24 | OG0008783 | Q94KE2 (TIC_ARATH) | Protein TIME FOR COFFEE | depleted | -1.14 | -3.91 | 9.3e-05 | 0.0001 | 0.0264 | 0.348 | 0.348 |
| 25 | OG0010059 | Q9LVJ1 (SBT14_ARATH) | Subtilisin-like protease SBT1.4 | depleted | -1.08 | -3.88 | 0.000106 | 0.0001 | 0.0264 | 0.135 | 0.135 |
| 26 | OG0006954 | F4IV99 (CHR5_ARATH) | Protein CHROMATIN REMODELING 5 | enriched | 1.04 | 3.87 | 0.000108 | 0.0001 | 0.0264 | 0.865 | 0.865 |
| 27 | OG0000163 | F4IIM1 (CSI1_ARATH) | Protein CELLULOSE SYNTHASE INTERACTIVE 1 | enriched | 1.06 | 3.87 | 0.00011 | 0.0001 | 0.0264 | 0.865 | 0.865 |
| 28 | OG0010177 | Q9M9C5 (Y1680_ARATH) | Probable leucine-rich repeat receptor-like protein kinase At1g68400 | depleted | -1.09 | -3.86 | 0.000112 | 0.0001 | 0.0264 | 0.135 | 0.135 |
| 29 | OG0000059 | F4HZB2 (BCHA1_ARATH) | Protein SPIRRIG | enriched | 0.871 | 3.84 | 0.000125 | 0.0001 | 0.0264 | 0.888 | 0.888 |
| 30 | OG0009680 | Q9SLI8 (2AB2D_ARATH) | Probable serine/threonine protein phosphatase 2A regulatory subunit B"delta | depleted | -1.08 | -3.81 | 0.000141 | 0.0001 | 0.0264 | 0.135 | 0.135 |
| 31 | OG0008764 | P0CB16 (DRL25_ARATH) | Putative disease resistance protein At4g19050 | depleted | -1.03 | -3.79 | 0.000153 | 0.0001 | 0.0264 | 0.18 | 0.18 |
| 32 | OG0007891 | Q7X9V2 (PIE1_ARATH) | Protein PHOTOPERIOD-INDEPENDENT EARLY FLOWERING 1 | enriched | 0.893 | 3.77 | 0.000165 | 0.0001 | 0.0264 | 0.888 | 0.888 |
| 33 | OG0009796 | C0LGP4 (Y3475_ARATH) | Probable LRR receptor-like serine/threonine-protein kinase At3g47570 | depleted | -0.902 | -3.53 | 0.00041 | 0.0001 | 0.0264 | 0.157 | 0.157 |
| 34 | OG0008698 | A4GGA5 (RPOB_PHAVU) | DNA-directed RNA polymerase subunit beta | depleted | -0.879 | -3.49 | 0.000481 | 0.0001 | 0.0264 | 0.169 | 0.169 |
| 35 | OG0006605 | Q9SRU2 (BIG_ARATH) | Auxin transport protein BIG | depleted | -0.915 | -3.44 | 0.000582 | 0.0001 | 0.0264 | 0.169 | 0.169 |
| 36 | OG0008980 | P10978 (POLX_TOBAC) | Retrovirus-related Pol polyprotein from transposon TNT 1-94 | depleted | -0.93 | -3.44 | 0.000585 | 0.0001 | 0.0264 | 0.169 | 0.169 |
| 37 | OG0008138 | P92523 (M860_ARATH) | Uncharacterized mitochondrial protein AtMg00860 | depleted | -0.913 | -3.33 | 0.000858 | 0.0001 | 0.0264 | 0.157 | 0.157 |
| 38 | OG0000327 | O23372 (ATXR3_ARATH) | Histone-lysine N-methyltransferase ATXR3 | enriched | 0.761 | 3.33 | 0.000884 | 0.0001 | 0.0264 | 0.91 | 0.91 |
| 39 | OG0006810 | Q9FFF9 (APC1_ARATH) | Anaphase-promoting complex subunit 1 | enriched | 0.732 | 3.31 | 0.000918 | 0.0001 | 0.0264 | 0.91 | 0.91 |
| 40 | OG0001995 | P31843 (RRPO_OENBE) | RNA-directed DNA polymerase homolog | enriched | 0.762 | 3.3 | 0.000975 | 0.0001 | 0.0264 | 0.91 | 0.91 |
| 41 | OG0012180 | Q8VY27 (E70H1_ARATH) | Exocyst complex component EXO70H1 | depleted | -0.742 | -3.24 | 0.00117 | 0.0001 | 0.0264 | 0.0899 | 0.0899 |
| 42 | OG0012763 | Q94HW2 (POLR1_ARATH) | Retrovirus-related Pol polyprotein from transposon RE1 | depleted | -0.737 | -3.23 | 0.00124 | 0.0001 | 0.0264 | 0.0899 | 0.0899 |
| 43 | OG0007467 | Q94HW2 (POLR1_ARATH) | Retrovirus-related Pol polyprotein from transposon RE1 | enriched | 0.729 | 3.22 | 0.00128 | 0.0001 | 0.0264 | 0.921 | 0.921 |
| 44 | OG0009300 | F4IAT2 (THOC2_ARATH) | THO complex subunit 2 | depleted | -0.749 | -3.17 | 0.00152 | 0.0001 | 0.0264 | 0.157 | 0.157 |
| 45 | OG0014463 | P31843 (RRPO_OENBE) | RNA-directed DNA polymerase homolog | depleted | -0.701 | -3.14 | 0.00168 | 0.0001 | 0.0264 | 0.0674 | 0.0674 |
| 46 | OG0010524 | Q9C8E7 (GLR33_ARATH) | Glutamate receptor 3.3 | depleted | -0.738 | -3.12 | 0.00182 | 0.0001 | 0.0264 | 0.0899 | 0.0899 |
| 47 | OG0012178 | P93050 (RKF3_ARATH) | Probable LRR receptor-like serine/threonine-protein kinase RKF3 | depleted | -0.74 | -3.11 | 0.00188 | 0.0001 | 0.0264 | 0.0899 | 0.0899 |
| 48 | OG0010178 | O80560 (IP5PC_ARATH) | Type I inositol polyphosphate 5-phosphatase 12 | depleted | -0.855 | -3.09 | 0.00202 | 0.0001 | 0.0264 | 0.157 | 0.157 |
| 49 | OG0006918 | F4J7T3 (EAF1A_ARATH) | Chromatin modification-related protein EAF1 A | enriched | 0.701 | 3.07 | 0.00213 | 0.0001 | 0.0264 | 0.921 | 0.921 |
| 50 | OG0010505 | O04408 (KSA_PEA) | Ent-copalyl diphosphate synthase, chloroplastic | depleted | -0.68 | -3 | 0.00269 | 0.0001 | 0.0264 | 0.0899 | 0.0899 |

#### Supplementary Table S5

Top 50 ranked copy-number association hits for marine lifestyle across 106 mammalian species. Rows correspond to orthogroups and include rank, orthogroup identifier, UniProt hit, putative function, effect direction, effect estimate, test statistic, raw p value, calibrated p value, false-discovery rate, prevalence, and mean copy number.

| Rank | Orthogroup | UniProt | Putative_function | Direction | Beta | z | raw_p | cal_p | FDR | Prev | Mean_CN |
|---|---|---|---|---|---|---|---|---|---|---|---|
| 1 | OG0018662 | Q9NYW5 (TA2R4_HUMAN) | Taste receptor type 2 member 4 | depleted | -1.93 | -7.34 | 2.15e-13 | 1.43e-05 | 0.00861 | 0.66 | 0.66 |
| 2 | OG0005961 | Q95MP7 (FRIH_CANLF) | Ferritin heavy chain | enriched | 1.69 | 6.57 | 5.04e-11 | 1.43e-05 | 0.00861 | 0.396 | 1.4 |
| 3 | OG0018317 | Q9H342 (O51J1_HUMAN) | Olfactory receptor 51J1 | depleted | -1.73 | -6.54 | 6.07e-11 | 1.43e-05 | 0.00861 | 0.67 | 0.708 |
| 4 | OG0004577 | Q9H211 (CDT1_HUMAN) | DNA replication factor Cdt1 | enriched | 1.67 | 6.35 | 2.12e-10 | 1.43e-05 | 0.00861 | 0.991 | 1.63 |
| 5 | OG0023673 | P00829 (ATPB_BOVIN) | ATP synthase F(1) complex catalytic subunit beta, mitochondrial | enriched | 1.75 | 6.19 | 5.91e-10 | 1.43e-05 | 0.00861 | 0.245 | 0.245 |
| 6 | OG0023699 | P53007 (TXTP_HUMAN) | Tricarboxylate transport protein, mitochondrial | enriched | 1.75 | 6.19 | 5.91e-10 | 1.43e-05 | 0.00861 | 0.245 | 0.245 |
| 7 | OG0023738 | Q3SWZ6 (SBDS_BOVIN) | Ribosome maturation protein SBDS | enriched | 1.75 | 6.19 | 5.91e-10 | 1.43e-05 | 0.00861 | 0.245 | 0.245 |
| 8 | OG0018208 | A8MTJ3 (GNAT3_HUMAN) | Guanine nucleotide-binding protein G(t) subunit alpha-3 | depleted | -1.67 | -6.16 | 7.07e-10 | 1.43e-05 | 0.00861 | 0.726 | 0.726 |
| 9 | OG0002665 | Q5Y4Z0 (T2R62_PANPA) | Taste receptor type 2 member 62 | enriched | 1.61 | 6.13 | 8.95e-10 | 1.43e-05 | 0.00861 | 0.632 | 1.99 |
| 10 | OG0020214 | Q3SZL8 (CNDH2_BOVIN) | Condensin-2 complex subunit H2 | enriched | 1.68 | 6.04 | 1.55e-09 | 1.43e-05 | 0.00861 | 0.245 | 0.481 |
| 11 | OG0018181 | Q148H8 (K2C72_BOVIN) | Keratin, type II cytoskeletal 72 | depleted | -1.68 | -6.04 | 1.58e-09 | 1.43e-05 | 0.00861 | 0.726 | 0.726 |
| 12 | OG0020938 | Q8NGY7 (O10J6_HUMAN) | Putative olfactory receptor 10J6 | enriched | 1.55 | 6.03 | 1.69e-09 | 1.43e-05 | 0.00861 | 0.358 | 0.406 |
| 13 | OG0022127 | A3FKF7 (G3P_MUSPF) | Glyceraldehyde-3-phosphate dehydrogenase | enriched | 1.58 | 6 | 1.93e-09 | 1.43e-05 | 0.00861 | 0.321 | 0.321 |
| 14 | OG0024002 | Q92733 (PRCC_HUMAN) | Proline-rich protein PRCC | enriched | 1.67 | 6 | 1.96e-09 | 1.43e-05 | 0.00861 | 0.236 | 0.236 |
| 15 | OG0024268 | Q9C0B7 (TNG6_HUMAN) | Transport and Golgi organization protein 6 homolog | enriched | 1.65 | 5.95 | 2.74e-09 | 1.43e-05 | 0.00861 | 0.226 | 0.226 |
| 16 | OG0024276 | Q69ZQ2 (ISY1_MOUSE) | Pre-mRNA-splicing factor ISY1 homolog | enriched | 1.65 | 5.95 | 2.74e-09 | 1.43e-05 | 0.00861 | 0.226 | 0.226 |
| 17 | OG0024287 | O95985 (TOP3B_HUMAN) | DNA topoisomerase 3-beta-1 | enriched | 1.65 | 5.95 | 2.74e-09 | 1.43e-05 | 0.00861 | 0.226 | 0.226 |
| 18 | OG0018505 | Q8NH63 (O51H1_HUMAN) | Olfactory receptor 51H1 | depleted | -1.56 | -5.95 | 2.75e-09 | 1.43e-05 | 0.00861 | 0.613 | 0.679 |
| 19 | OG0018663 | Q646G9 (TA2R1_PANPA) | Taste receptor type 2 member 1 | depleted | -1.57 | -5.91 | 3.32e-09 | 1.43e-05 | 0.00861 | 0.651 | 0.66 |
| 20 | OG0018023 | B0LKP1 (KRT35_SHEEP) | Keratin, type I cuticular Ha5 | depleted | -1.64 | -5.91 | 3.43e-09 | 1.43e-05 | 0.00861 | 0.736 | 0.745 |
| 21 | OG0023529 | Q865K9 (MPRB_PIG) | Membrane progestin receptor beta | enriched | 1.66 | 5.9 | 3.69e-09 | 1.43e-05 | 0.00861 | 0.255 | 0.255 |
| 22 | OG0023958 | Q9HCJ2 (LRC4C_HUMAN) | Leucine-rich repeat-containing protein 4C | enriched | 1.63 | 5.9 | 3.72e-09 | 1.43e-05 | 0.00861 | 0.236 | 0.236 |
| 23 | OG0023935 | Q9Y490 (TLN1_HUMAN) | Talin-1 | enriched | 1.63 | 5.88 | 4e-09 | 1.43e-05 | 0.00861 | 0.236 | 0.236 |
| 24 | OG0023973 | O70194 (EIF3D_MOUSE) | Eukaryotic translation initiation factor 3 subunit D | enriched | 1.63 | 5.88 | 4e-09 | 1.43e-05 | 0.00861 | 0.236 | 0.236 |
| 25 | OG0019940 | Q96NN9 (AIFM3_HUMAN) | Apoptosis-inducing factor 3 | enriched | 1.67 | 5.88 | 4.09e-09 | 1.43e-05 | 0.00861 | 0.255 | 0.509 |
| 26 | OG0018170 | Q8NH61 (O51F2_HUMAN) | Olfactory receptor 51F2 | depleted | -1.44 | -5.88 | 4.15e-09 | 1.43e-05 | 0.00861 | 0.642 | 0.726 |
| 27 | OG0006413 | Q6IEU7 (OR5MA_HUMAN) | Olfactory receptor 5M10 | depleted | -1.42 | -5.86 | 4.66e-09 | 1.43e-05 | 0.00861 | 0.698 | 1.34 |
| 28 | OG0024451 | O18965 (KCNH1_BOVIN) | Voltage-gated delayed rectifier potassium channel KCNH1 | enriched | 1.62 | 5.86 | 4.74e-09 | 1.43e-05 | 0.00861 | 0.217 | 0.217 |
| 29 | OG0024454 | Q9H165 (BC11A_HUMAN) | BCL11 transcription factor A | enriched | 1.62 | 5.86 | 4.74e-09 | 1.43e-05 | 0.00861 | 0.217 | 0.217 |
| 30 | OG0024484 | F1N6G5 (HACE1_BOVIN) | E3 ubiquitin-protein ligase HACE1 | enriched | 1.62 | 5.86 | 4.74e-09 | 1.43e-05 | 0.00861 | 0.217 | 0.217 |
| 31 | OG0024527 | Q06AK6 (TFP11_PIG) | Tuftelin-interacting protein 11 | enriched | 1.62 | 5.86 | 4.74e-09 | 1.43e-05 | 0.00861 | 0.217 | 0.217 |
| 32 | OG0017440 | P97434 (MPRIP_MOUSE) | Myosin phosphatase Rho-interacting protein | enriched | 1.6 | 5.84 | 5.08e-09 | 1.43e-05 | 0.00861 | 0.236 | 0.821 |
| 33 | OG0019187 | Q9NY15 (STAB1_HUMAN) | Stabilin-1 | enriched | 1.61 | 5.84 | 5.25e-09 | 1.43e-05 | 0.00861 | 0.255 | 0.594 |
| 34 | OG0023594 | P07195 (LDHB_HUMAN) | L-lactate dehydrogenase B chain | enriched | 1.6 | 5.79 | 7.11e-09 | 1.43e-05 | 0.00861 | 0.236 | 0.245 |
| 35 | OG0019779 | P60990 (PIP_RABIT) | Prolactin-inducible protein homolog | depleted | -1.56 | -5.78 | 7.26e-09 | 1.43e-05 | 0.00861 | 0.528 | 0.528 |
| 36 | OG0004128 | P54098 (DPOG1_HUMAN) | DNA polymerase subunit gamma-1 | enriched | 1.6 | 5.77 | 7.95e-09 | 1.43e-05 | 0.00861 | 1 | 1.73 |
| 37 | OG0018652 | Q8N127 (O5AS1_HUMAN) | Olfactory receptor 5AS1 | depleted | -1.5 | -5.76 | 8.27e-09 | 1.43e-05 | 0.00861 | 0.66 | 0.66 |
| 38 | OG0018072 | Q8NGK1 (O51G1_HUMAN) | Olfactory receptor 51G1 | depleted | -1.46 | -5.76 | 8.55e-09 | 1.43e-05 | 0.00861 | 0.66 | 0.736 |
| 39 | OG0024179 | Q9NQW6 (ANLN_HUMAN) | Anillin | enriched | 1.57 | 5.76 | 8.56e-09 | 1.43e-05 | 0.00861 | 0.226 | 0.226 |
| 40 | OG0024216 | A5D7P8 (SOSB2_BOVIN) | SOSS complex subunit B2 | enriched | 1.57 | 5.76 | 8.56e-09 | 1.43e-05 | 0.00861 | 0.226 | 0.226 |
| 41 | OG0018870 | Q8NH63 (O51H1_HUMAN) | Olfactory receptor 51H1 | depleted | -1.5 | -5.74 | 9.21e-09 | 1.43e-05 | 0.00861 | 0.604 | 0.632 |
| 42 | OG0024284 | Q5E9R1 (S52A3_BOVIN) | Solute carrier family 52, riboflavin transporter, member 3 | enriched | 1.56 | 5.74 | 9.31e-09 | 1.43e-05 | 0.00861 | 0.226 | 0.226 |
| 43 | OG0023658 | Q9UPR3 (SMG5_HUMAN) | Nonsense-mediated mRNA decay factor SMG5 | enriched | 1.59 | 5.74 | 9.59e-09 | 1.43e-05 | 0.00861 | 0.236 | 0.245 |
| 44 | OG0023697 | Q5BIS9 (AAKB1_BOVIN) | 5'-AMP-activated protein kinase subunit beta-1 | enriched | 1.6 | 5.74 | 9.7e-09 | 1.43e-05 | 0.00861 | 0.245 | 0.245 |
| 45 | OG0024175 | Q5SW79 (CE170_HUMAN) | Centrosomal protein of 170 kDa | enriched | 1.56 | 5.73 | 9.86e-09 | 1.43e-05 | 0.00861 | 0.226 | 0.226 |
| 46 | OG0018320 | Q8BFZ3 (ACTBL_MOUSE) | Beta-actin-like protein 2 | depleted | -1.58 | -5.71 | 1.15e-08 | 1.43e-05 | 0.00861 | 0.708 | 0.708 |
| 47 | OG0018825 | Q9H340 (O51B6_HUMAN) | Olfactory receptor 51B6 | depleted | -1.47 | -5.7 | 1.2e-08 | 1.43e-05 | 0.00861 | 0.557 | 0.642 |
| 48 | OG0023853 | P51977 (AL1A1_SHEEP) | Aldehyde dehydrogenase 1A1 | enriched | 1.54 | 5.69 | 1.24e-08 | 1.43e-05 | 0.00861 | 0.226 | 0.236 |
| 49 | OG0017815 | Q5SZD4 (GLYL3_HUMAN) | Glycine N-acyltransferase-like protein 3 | depleted | -1.55 | -5.68 | 1.36e-08 | 1.43e-05 | 0.00861 | 0.745 | 0.774 |
| 50 | OG0024534 | Q3ZBF0 (NADE_BOVIN) | Glutamine-dependent NAD(+) synthetase | enriched | 1.54 | 5.67 | 1.41e-08 | 1.43e-05 | 0.00861 | 0.217 | 0.217 |

#### Supplementary Table S6

Top 50 ranked presence/absence association hits for marine lifestyle across 106 mammalian species. Rows correspond to orthogroups and include rank, orthogroup identifier, UniProt hit, putative function, effect direction, effect estimate, test statistic, raw p value, calibrated p value, false-discovery rate, prevalence, and mean copy number.

| Rank | Orthogroup | UniProt | Putative_function | Direction | Beta | z | raw_p | cal_p | FDR | Prev | Mean_CN |
|---|---|---|---|---|---|---|---|---|---|---|---|
| 1 | OG0018662 | Q9NYW5 (TA2R4_HUMAN) | Taste receptor type 2 member 4 | depleted | -1.93 | -7.34 | 2.15e-13 | 0.0001 | 0.00861 | 0.66 | 0.66 |
| 2 | OG0018317 | Q9H342 (O51J1_HUMAN) | Olfactory receptor 51J1 | depleted | -1.87 | -7.16 | 8.14e-13 | 0.0001 | 0.00861 | 0.67 | 0.67 |
| 3 | OG0018172 | Q9H343 (O51I1_HUMAN) | Olfactory receptor 51I1 | depleted | -1.81 | -7.04 | 1.94e-12 | 0.0001 | 0.00861 | 0.613 | 0.613 |
| 4 | OG0018170 | Q8NH61 (O51F2_HUMAN) | Olfactory receptor 51F2 | depleted | -1.81 | -7.02 | 2.29e-12 | 0.0001 | 0.00861 | 0.642 | 0.642 |
| 5 | OG0017497 | P34985 (OL143_MOUSE) | Olfactory receptor 8C8 | depleted | -1.79 | -6.93 | 4.08e-12 | 0.0001 | 0.00861 | 0.613 | 0.613 |
| 6 | OG0006413 | Q6IEU7 (OR5MA_HUMAN) | Olfactory receptor 5M10 | depleted | -1.82 | -6.92 | 4.59e-12 | 0.0001 | 0.00861 | 0.698 | 0.698 |
| 7 | OG0017494 | Q8NGJ5 (O51L1_HUMAN) | Olfactory receptor 51L1 | depleted | -1.84 | -6.79 | 1.13e-11 | 0.0001 | 0.00861 | 0.726 | 0.726 |
| 8 | OG0017813 | Q5QD14 (TAAR5_MOUSE) | Trace amine-associated receptor 5 | depleted | -1.74 | -6.58 | 4.68e-11 | 0.0001 | 0.00861 | 0.708 | 0.708 |
| 9 | OG0005961 | Q95MP7 (FRIH_CANLF) | Ferritin heavy chain | enriched | 1.82 | 6.5 | 8.14e-11 | 0.0001 | 0.00861 | 0.396 | 0.396 |
| 10 | OG0007700 | A0A2R8Y4L6 (OR5D3_HUMAN) | Olfactory receptor 5D3 | depleted | -1.64 | -6.48 | 9.38e-11 | 0.0001 | 0.00861 | 0.547 | 0.547 |
| 11 | OG0018825 | Q9H340 (O51B6_HUMAN) | Olfactory receptor 51B6 | depleted | -1.7 | -6.45 | 1.09e-10 | 0.0001 | 0.00861 | 0.557 | 0.557 |
| 12 | OG0018512 | Q8NGW6 (OR6K6_HUMAN) | Olfactory receptor 6K6 | depleted | -1.67 | -6.43 | 1.24e-10 | 0.0001 | 0.00861 | 0.613 | 0.613 |
| 13 | OG0018505 | Q8NH63 (O51H1_HUMAN) | Olfactory receptor 51H1 | depleted | -1.66 | -6.43 | 1.26e-10 | 0.0001 | 0.00861 | 0.613 | 0.613 |
| 14 | OG0020938 | Q8NGY7 (O10J6_HUMAN) | Putative olfactory receptor 10J6 | enriched | 1.63 | 6.36 | 2.02e-10 | 0.0001 | 0.00861 | 0.358 | 0.358 |
| 15 | OG0017806 | Q8NGT1 (OR2K2_HUMAN) | Olfactory receptor 2K2 | depleted | -1.77 | -6.35 | 2.09e-10 | 0.0001 | 0.00861 | 0.736 | 0.736 |
| 16 | OG0002175 | Q8NGF9 (OR4X2_HUMAN) | Olfactory receptor 4X2 | depleted | -1.65 | -6.34 | 2.32e-10 | 0.0001 | 0.00861 | 0.585 | 0.585 |
| 17 | OG0017092 | Q6EIZ0 (K1C10_CANLF) | Keratin, type I cytoskeletal 10 | depleted | -1.75 | -6.26 | 3.8e-10 | 0.0001 | 0.00861 | 0.736 | 0.736 |
| 18 | OG0006417 | Q8NH73 (OR4S2_HUMAN) | Olfactory receptor 4S2 | depleted | -1.62 | -6.24 | 4.35e-10 | 0.0001 | 0.00861 | 0.642 | 0.642 |
| 19 | OG0018653 | Q8NGK4 (O52K1_HUMAN) | Olfactory receptor 52K1 | depleted | -1.59 | -6.23 | 4.67e-10 | 0.0001 | 0.00861 | 0.528 | 0.528 |
| 20 | OG0017658 | Q8NGJ2 (O52H1_HUMAN) | Olfactory receptor 52H1 | depleted | -1.59 | -6.2 | 5.55e-10 | 0.0001 | 0.00861 | 0.538 | 0.538 |
| 21 | OG0006258 | Q9H346 (O52D1_HUMAN) | Olfactory receptor 52D1 | depleted | -1.75 | -6.19 | 5.91e-10 | 0.0001 | 0.00861 | 0.755 | 0.755 |
| 22 | OG0015421 | Q6A152 (CP4X1_MOUSE) | Cytochrome P450 4X1 | depleted | -1.75 | -6.19 | 5.91e-10 | 0.0001 | 0.00861 | 0.755 | 0.755 |
| 23 | OG0017093 | Q6EIY9 (K2C1_CANLF) | Keratin, type II cytoskeletal 1 | depleted | -1.75 | -6.19 | 5.91e-10 | 0.0001 | 0.00861 | 0.755 | 0.755 |
| 24 | OG0017646 | O75635 (SPB7_HUMAN) | Serpin B7 | depleted | -1.75 | -6.19 | 5.91e-10 | 0.0001 | 0.00861 | 0.755 | 0.755 |
| 25 | OG0020214 | Q3SZL8 (CNDH2_BOVIN) | Condensin-2 complex subunit H2 | enriched | 1.75 | 6.19 | 5.91e-10 | 0.0001 | 0.00861 | 0.245 | 0.245 |
| 26 | OG0021530 | Q99627 (CSN8_HUMAN) | COP9 signalosome complex subunit 8 | enriched | 1.75 | 6.19 | 5.91e-10 | 0.0001 | 0.00861 | 0.245 | 0.245 |
| 27 | OG0023673 | P00829 (ATPB_BOVIN) | ATP synthase F(1) complex catalytic subunit beta, mitochondrial | enriched | 1.75 | 6.19 | 5.91e-10 | 0.0001 | 0.00861 | 0.245 | 0.245 |
| 28 | OG0023699 | P53007 (TXTP_HUMAN) | Tricarboxylate transport protein, mitochondrial | enriched | 1.75 | 6.19 | 5.91e-10 | 0.0001 | 0.00861 | 0.245 | 0.245 |
| 29 | OG0023738 | Q3SWZ6 (SBDS_BOVIN) | Ribosome maturation protein SBDS | enriched | 1.75 | 6.19 | 5.91e-10 | 0.0001 | 0.00861 | 0.245 | 0.245 |
| 30 | OG0018073 | Q8NGK0 (O51G2_HUMAN) | Olfactory receptor 51G2 | depleted | -1.57 | -6.18 | 6.35e-10 | 0.0001 | 0.00861 | 0.585 | 0.585 |
| 31 | OG0018208 | A8MTJ3 (GNAT3_HUMAN) | Guanine nucleotide-binding protein G(t) subunit alpha-3 | depleted | -1.67 | -6.16 | 7.07e-10 | 0.0001 | 0.00861 | 0.726 | 0.726 |
| 32 | OG0019103 | Q6IWH7 (ANO7_HUMAN) | Anoctamin-7 | enriched | 1.6 | 6.16 | 7.27e-10 | 0.0001 | 0.00861 | 0.425 | 0.425 |
| 33 | OG0017007 | Q8NGK0 (O51G2_HUMAN) | Olfactory receptor 51G2 | depleted | -1.59 | -6.14 | 8.04e-10 | 0.0001 | 0.00861 | 0.623 | 0.623 |
| 34 | OG0015656 | Q8NGP3 (OR5M9_HUMAN) | Olfactory receptor 5M9 | depleted | -1.55 | -6.14 | 8.07e-10 | 0.0001 | 0.00861 | 0.557 | 0.557 |
| 35 | OG0018582 | Q8VEZ1 (OR4CC_MOUSE) | Olfactory receptor 4C12 | depleted | -1.55 | -6.12 | 9.4e-10 | 0.0001 | 0.00861 | 0.557 | 0.557 |
| 36 | OG0003549 | Q8NH72 (OR4C6_HUMAN) | Olfactory receptor 4C6 | depleted | -1.56 | -6.11 | 1.01e-09 | 0.0001 | 0.00861 | 0.613 | 0.613 |
| 37 | OG0003718 | Q96JA4 (M4A14_HUMAN) | Membrane-spanning 4-domains subfamily A member 14 | depleted | -1.68 | -6.05 | 1.48e-09 | 0.0001 | 0.00861 | 0.726 | 0.726 |
| 38 | OG0023594 | P07195 (LDHB_HUMAN) | L-lactate dehydrogenase B chain | enriched | 1.68 | 6.04 | 1.54e-09 | 0.0001 | 0.00861 | 0.236 | 0.236 |
| 39 | OG0018181 | Q148H8 (K2C72_BOVIN) | Keratin, type II cytoskeletal 72 | depleted | -1.68 | -6.04 | 1.58e-09 | 0.0001 | 0.00861 | 0.726 | 0.726 |
| 40 | OG0014719 | P22760 (AAAD_HUMAN) | Arylacetamide deacetylase | depleted | -1.62 | -6.03 | 1.68e-09 | 0.0001 | 0.00861 | 0.726 | 0.726 |
| 41 | OG0018318 | Q6IF00 (OR2T2_HUMAN) | Olfactory receptor 2T2 | depleted | -1.57 | -6.01 | 1.84e-09 | 0.0001 | 0.00861 | 0.481 | 0.481 |
| 42 | OG0018023 | B0LKP1 (KRT35_SHEEP) | Keratin, type I cuticular Ha5 | depleted | -1.66 | -6.01 | 1.85e-09 | 0.0001 | 0.00861 | 0.736 | 0.736 |
| 43 | OG0022127 | A3FKF7 (G3P_MUSPF) | Glyceraldehyde-3-phosphate dehydrogenase | enriched | 1.58 | 6 | 1.93e-09 | 0.0001 | 0.00861 | 0.321 | 0.321 |
| 44 | OG0017440 | P97434 (MPRIP_MOUSE) | Myosin phosphatase Rho-interacting protein | enriched | 1.67 | 6 | 1.96e-09 | 0.0001 | 0.00861 | 0.236 | 0.236 |
| 45 | OG0022084 | O00562 (PITM1_HUMAN) | Membrane-associated phosphatidylinositol transfer protein 1 | enriched | 1.67 | 6 | 1.96e-09 | 0.0001 | 0.00861 | 0.236 | 0.236 |
| 46 | OG0024002 | Q92733 (PRCC_HUMAN) | Proline-rich protein PRCC | enriched | 1.67 | 6 | 1.96e-09 | 0.0001 | 0.00861 | 0.236 | 0.236 |
| 47 | OG0002383 | P00178 (CP2B4_RABIT) | Cytochrome P450 2B4 | depleted | -1.68 | -5.98 | 2.23e-09 | 0.0001 | 0.00861 | 0.745 | 0.745 |
| 48 | OG0002823 | Q8NH60 (O52J3_HUMAN) | Olfactory receptor 52J3 | depleted | -1.68 | -5.98 | 2.23e-09 | 0.0001 | 0.00861 | 0.745 | 0.745 |
| 49 | OG0016394 | Q8NH37 (OR4C3_HUMAN) | Olfactory receptor 4C3 | depleted | -1.54 | -5.96 | 2.53e-09 | 0.0001 | 0.00861 | 0.594 | 0.594 |
| 50 | OG0023658 | Q9UPR3 (SMG5_HUMAN) | Nonsense-mediated mRNA decay factor SMG5 | enriched | 1.65 | 5.96 | 2.54e-09 | 0.0001 | 0.00861 | 0.236 | 0.236 |

#### Supplementary Table S7

Complete bacterial species list for the diazotrophy analysis, including phenotype labels and genome-quality metadata used in the OrthoGLMM model.

| Species | diazotrophy | BUSCO |
|---|---|---|
| Abiotrophia_defectiva | 0 | 98.4 |
| Acanthopleuribacter_pedis | 0 | 96.0 |
| Acetanaerobacterium_elongatum | 0 | 94.4 |
| Acetitomaculum_ruminis_DSM_5522 | 0 | 99.2 |
| Acetivibrio_straminisolvens | 0 | 99.2 |
| Acetoanaerobium_noterae | 0 | 98.4 |
| Acetobacter_farinalis | 0 | 100.0 |
| Acetobacterium_bakii | 0 | 99.2 |
| Acetohalobium_arabaticum_DSM_5501 | 0 | 99.2 |
| Acetomicrobium_thermoterrenum_DSM_13490 | 0 | 90.3 |
| Acetonema_longum_DSM_6540 | 0 | 98.4 |
| Acholeplasma_equirhinis | 0 | 82.3 |
| Achromobacter_anxifer | 0 | 99.2 |
| Acidaminobacter_hydrogenoformans_DSM_2784 | 0 | 99.2 |
| Acidaminococcus_massiliensis | 0 | 96.8 |
| Acidianus_infernus | 0 | 14.5 |
| Acidicaldus_organivorans | 0 | 79.0 |
| Acidicapsa_dinghuensis | 0 | 96.8 |
| Acidiferrobacter_thiooxydans | 0 | 97.6 |
| Acidimicrobium_ferrooxidans_DSM_10331 | 0 | 92.7 |
| Acidiphilium_rubrum | 0 | 98.4 |
| Acidiplasma_aeolicum | 0 | 14.5 |
| Acidisoma_cellulosilyticum | 0 | 97.6 |
| Acidisphaera_rubrifaciens_HS-AP3 | 0 | 86.3 |
| Aciditerrimonas_ferrireducens | 0 | 92.7 |
| Acidithiobacillus_albertensis | 0 | 98.4 |
| Acidobacterium_capsulatum_ATCC_51196 | 0 | 95.2 |
| Acidocella_aquatica | 0 | 99.2 |
| Acidomonas_methanolica | 0 | 99.2 |
| Acidothermus_cellulolyticus_11B | 0 | 98.4 |
| Acidovorax_lacteus | 0 | 100.0 |
| Acinetobacter_amyesii | 0 | 99.2 |
| Acinetobacter_apis | 0 | 100.0 |
| Acrocarpospora_corrugata | 0 | 100.0 |
| Actinoallomurus_liliacearum | 0 | 100.0 |
| Actinobacillus_capsulatus_DSM_19761 | 0 | 99.2 |
| Actinobaculum_massiliense_ACS-171-V-Col2 | 0 | 95.2 |
| Actinocatenispora_comari | 0 | 98.4 |
| Actinocorallia_longicatena | 0 | 99.2 |
| Actinomadura_geliboluensis | 0 | 100.0 |
| Actinomyces_lilanjuaniae | 0 | 84.7 |
| Actinophytocola_glycyrrhizae | 0 | 99.2 |
| Actinoplanes_hulinensis | 0 | 99.2 |
| Actinopolymorpha_singaporensis | 0 | 99.2 |
| Actinopolyspora_biskrensis | 0 | 100.0 |
| Actinospica_robiniae_DSM_44927 | 0 | 98.4 |
| Actinosynnema_pretiosum | 0 | 100.0 |
| Actinotalea_lenta | 0 | 100.0 |
| Adhaeribacter_rhizoryzae | 0 | 97.6 |
| Aequorivita_viscosa | 0 | 98.4 |
| Aeriscardovia_aeriphila | 0 | 88.7 |
| Aerococcus_kribbianus | 0 | 90.3 |
| Aeromicrobium_choanae | 0 | 97.6 |
| Aeromonas_aquatica | 0 | 98.4 |
| Aeropyrum_camini_SY1___JCM_12091 | 0 | 12.9 |
| Aestuariimicrobium_ganziense | 0 | 98.4 |
| Afipia_birgiae_34632 | 0 | 97.6 |
| Agreia_pratensis | 0 | 97.6 |
| Agrobacterium_cavarae | 0 | 100.0 |
| Agrococcus_phoenicis | 0 | 96.8 |
| Agromyces_intestinalis | 0 | 96.8 |
| Akkermansia_massiliensis | 0 | 83.9 |
| Albimonas_pacifica | 0 | 99.2 |

| Species | diazotrophy | BUSCO |
|---|---|---|
| Alcaligenes_endophyticus | 0 | 100.0 |
| Alcanivorax_limicola | 0 | 100.0 |
| Algibacter_amylolyticus | 0 | 100.0 |
| Algiphilus_aromaticivorans_DG1253 | 0 | 100.0 |
| Algisphaera_agarilytica | 0 | 87.1 |
| Algoriphagus_machipongonensis | 0 | 97.6 |
| Alicycliphilus_soli | 0 | 100.0 |
| Alicyclobacillus_fodiniaquatilis | 0 | 100.0 |
| Aliivibrio_finisterrensis | 0 | 100.0 |
| Alishewanella_longhuensis | 0 | 98.4 |
| Alistipes_finegoldii | 0 | 93.5 |
| Alkalibacillus_silvisoli | 0 | 100.0 |
| Alkalibacter_saccharofermentans_DSM_14828 | 0 | 99.2 |
| Alkalibacterium_indicireducens | 0 | 97.6 |
| Alkalibaculum_bacchi | 0 | 97.6 |
| Alkaliflexus_imshenetskii_DSM_15055 | 0 | 93.5 |
| Alkalimarinus_sediminis | 0 | 100.0 |
| Alkaliphilus_peptidifermentans_DSM_18978 | 0 | 100.0 |
| Alkalitalea_saponilacus | 0 | 100.0 |
| Alkanindiges_illinoisensis_DSM_15370 | 0 | 100.0 |
| Allobaculum_stercoricanis_DSM_13633 | 0 | 94.4 |
| Allocatelliglobosispora_scoriae | 0 | 98.4 |
| Allochromatium_warmingii | 1 | 97.6 |
| Alloprevotella_tannerae_ATCC_51259 | 0 | 98.4 |
| Allorhizobium_borbori | 1 | 100.0 |
| Alloscardovia_theropitheci | 0 | 94.4 |
| Alteromonas_arenosi | 0 | 100.0 |
| Amaricoccus_tamworthensis | 0 | 99.2 |
| Aminobacter_ciceronei | 0 | 99.2 |
| Aminobacterium_mobile_DSM_12262 | 0 | 89.5 |
| Aminomonas_paucivorans_DSM_12260 | 0 | 89.5 |
| Ammonifex_thiophilus | 0 | 99.2 |
| Ammoniphilus_resinae | 0 | 100.0 |
| Amnibacterium_endophyticum | 0 | 97.6 |
| Amorphus_coralli_DSM_19760 | 0 | 99.2 |
| Amphibacillus_indicireducens | 0 | 97.6 |
| Amycolatopsis_dongchuanensis | 0 | 98.4 |
| Anabaena_catenula_FACHB-362 | 1 | 96.8 |
| Anaeroarcus_burkinensis_DSM_6283 | 0 | 99.2 |
| Anaerobacillus_alkaliphilus | 1 | 100.0 |
| Anaerobiospirillum_succiniciproducens_DSM_6400 | 0 | 99.2 |
| Anaerocellum_danielii | 0 | 98.4 |
| Anaerococcus_cruorum | 0 | 90.3 |
| Anaerofilum_hominis | 0 | 98.4 |
| Anaerolinea_thermolimosa | 0 | 91.9 |
| Anaeromusa_acidaminophila_DSM_3853 | 0 | 99.2 |
| Anaeromyxobacter_soli | 0 | 96.0 |
| Anaerophaga_thermohalophila_DSM_12881 | 0 | 96.8 |
| Anaeroplasma_bactoclasticum | 0 | 91.9 |
| Anaerostipes_amylophilus | 0 | 99.2 |
| Anaerotruncus_rubiinfantis | 0 | 95.2 |
| Anaerovibrio_lipolyticus_DSM_3074 | 0 | 96.8 |
| Anaerovirgula_multivorans | 0 | 100.0 |
| Anaerovorax_odorimutans_DSM_5092 | 0 | 99.2 |
| Ancylobacter_defluvii | 0 | 99.2 |
| Andreprevotia_lacus_DSM_23236 | 0 | 100.0 |
| Aneurinibacillus_migulanus | 0 | 100.0 |
| Angustibacter_luteus | 0 | 97.6 |
| Anoxybacillus_eryuanensis | 0 | 100.0 |
| Anoxynatronum_buryatiense | 0 | 100.0 |
| Antarctobacter_heliothermus | 0 | 99.2 |
| Antricoccus_suffuscus | 0 | 97.6 |
| Aphanothece_microscopica_RSMan92 | 0 | 95.2 |
| Aquabacter_sediminis | 0 | 99.2 |
| Aquabacterium_humicola | 0 | 100.0 |
| Aquamicrobium_terrae | 0 | 100.0 |
| Aquaspirillum_serpens_DSM_68 | 0 | 100.0 |

| **Species** | **diazotrophy** | **BUSCO** |
|---|---|---|
| Aquifex_aeolicus_VF5 | 0 | 87.1 |
| Aquiflexum_balticum_DSM_16537 | 0 | 99.2 |
| Aquimarina_hainanensis | 0 | 98.4 |
| Aquincola_agrisoli | 0 | 98.4 |
| Aquipuribacter_hungaricus | 0 | 95.2 |
| Aquitalea_magnusonii | 0 | 100.0 |
| Arcanobacterium_phocae | 0 | 95.2 |
| Archaeoglobus_neptunius | 0 | 20.2 |
| Archangium_gephyra | 0 | 92.7 |
| Arcicella_aurantiaca | 0 | 89.5 |
| Arcobacter_peruensis | 1 | 91.1 |
| Arcticibacter_tournemirensis | 0 | 99.2 |
| Ardenticatena_maritima | 0 | 96.8 |
| Arenibacter_aquaticus | 0 | 100.0 |
| Arenicella_chitinivorans | 0 | 99.2 |
| Arenimonas_daejeonensis | 0 | 77.4 |
| Arhodomonas_aquaeolei_DSM_8974 | 0 | 100.0 |
| Arsenicicoccus_piscis | 0 | 98.4 |
| Arsenophonus_endosymbiont_of_Bemisia_tabaci_Q2 | 0 | 87.9 |
| Arthrobacter_antibioticus | 0 | 99.2 |
| Asticcacaulis_endophyticus | 0 | 98.4 |
| Atopobium_deltae | 0 | 92.7 |
| Atopostipes_suicloacalis_DSM_15692 | 0 | 92.7 |
| Aurantimonas_endophytica | 0 | 100.0 |
| Aureimonas_endophytica | 0 | 100.0 |
| Aureispira_anguillae | 0 | 96.8 |
| Aureitalea_marina | 0 | 98.4 |
| Azoarcus_indigens | 1 | 100.0 |
| Azospira_inquinata | 1 | 100.0 |
| Azospirillum_baldaniorum | 1 | 100.0 |
| Azotobacter_bryophylli | 1 | 99.2 |
| Azovibrio_restrictus_DSM_23866 | 1 | 98.4 |
| Bacteriovorax_antarcticus | 0 | 87.9 |
| Bacteroides_bouchesdurhonensis | 0 | 99.2 |
| Balneatrix_alpica_DSM_16621 | 0 | 99.2 |
| Balneola_vulgaris_DSM_17893 | 0 | 98.4 |
| Barnesiella_intestinihominis_YIT_11860 | 0 | 98.4 |
| Bauldia_litoralis | 0 | 99.2 |
| Beggiatoa_alba_B18LD | 1 | 98.4 |
| Beijerinckia_mobilis | 1 | 99.2 |
| Belliella_marina | 0 | 98.4 |
| Beutenbergia_cavernae_DSM_12333 | 0 | 97.6 |
| Bhargavaea_ullalensis | 0 | 100.0 |
| Bifidobacterium_actinocoloniiforme_DSM_22766 | 0 | 90.3 |
| Bilophila_wadsworthia | 0 | 93.5 |
| Bizionia_sediminis | 0 | 100.0 |
| Blastochloris_sulfoviridis | 0 | 100.0 |
| Blastococcus_deserti | 0 | 98.4 |
| Blastomonas_ursincola | 0 | 100.0 |
| Blastopirellula_sediminis | 0 | 85.5 |
| Blattabacterium_cuenoti | 0 | 78.2 |
| Blautia_ammoniilytica | 0 | 98.4 |
| Bordetella_ansorpii | 0 | 100.0 |
| Borrelia_crocidurae | 0 | 69.4 |
| Brachybacterium_equifaecis | 0 | 98.4 |
| Brachymonas_denitrificans_DSM_15123 | 0 | 99.2 |
| Brachyspira_aalborgi | 0 | 89.5 |
| Bradyrhizobium_canariense | 1 | 99.2 |
| Brenneria_corticis | 0 | 96.8 |
| Brevibacillus_antibioticus | 0 | 100.0 |
| Brevibacterium_album_DSM_18261 | 0 | 99.2 |
| Brevundimonas_alba | 0 | 100.0 |
| Brochothrix_campestris | 0 | 100.0 |
| Brooklawnia_cerclae | 0 | 97.6 |
| Brucella_abortus | 0 | 98.4 |
| Brumimicrobium_aurantiacum | 0 | 99.2 |
| Bryobacter_aggregatus_MPL3 | 0 | 94.4 |

| Species | diazotrophy | BUSCO |
|---|---|---|
| Bryocella_elongata | 0 | 92.7 |
| Budvicia_diplopodorum | 0 | 99.2 |
| Bulleidia_extructa_W1219 | 0 | 90.3 |
| Burkholderia_alba | 1 | 99.2 |
| Buttiauxella_massiliensis | 0 | 100.0 |
| Butyrivibrio_fibrisolvens | 0 | 97.6 |
| Byssovorax_cruenta_JCM_12614 | 0 | 29.8 |
| Caldanaerobacter_subterraneus_subsp._tengcongensis_MB4 | 0 | 100.0 |
| Caldibacillus_debilis_DSM_16016 | 0 | 96.0 |
| Caldicellulosiruptor_naganoensis | 0 | 98.4 |
| Caldicoprobacter_faecalis | 0 | 97.6 |
| Caldimicrobium_thiodismutans | 0 | 93.5 |
| Caldimonas_mangrovi | 0 | 100.0 |
| Calditerrivibrio_nitroreducens_DSM_19672 | 0 | 96.8 |
| Caldithrix_abyssi_DSM_13497 | 0 | 98.4 |
| Caldivirga_maquilingensis_IC-167 | 0 | 16.1 |
| Caloramator_quimbayensis | 0 | 96.8 |
| Caloranaerobacter_azorensis_DSM_13643 | 0 | 99.2 |
| Calothrix_parietina_FACHB-288 | 1 | 96.8 |
| Caminibacter_pacificus | 0 | 91.9 |
| Caminicella_sporogenes | 0 | 99.2 |
| Campylobacter_anatolicus | 0 | 92.7 |
| Capnocytophaga_bilenii | 0 | 100.0 |
| Carboxydothermus_islandicus | 0 | 98.4 |
| Cardiobacterium_valvarum | 0 | 87.1 |
| Carnobacterium_antarcticum | 0 | 96.0 |
| Catellatospora_aurea | 0 | 99.2 |
| Catellicoccus_marimammalium_M35_04_3 | 0 | 92.7 |
| Catenibacterium_faecis | 0 | 98.4 |
| Catenulispora_subtropica | 0 | 99.2 |
| Catenuloplanes_niger | 0 | 99.2 |
| Catonella_morbi_ATCC_51271 | 0 | 97.6 |
| Caulobacter_soli | 0 | 100.0 |
| Cecembia_rubra | 0 | 97.6 |
| Cedecea_selenatireducens | 0 | 98.4 |
| Celeribacter_arenosi | 0 | 96.8 |
| Cellulomonas_biazotea | 0 | 100.0 |
| Cellulophaga_tyrosinoxydans | 0 | 100.0 |
| Cellulosimicrobium_arenosum | 0 | 100.0 |
| Cellvibrio_chitinivorans | 1 | 100.0 |
| Cenarchaeum_symbiosum_A | 0 | 14.5 |
| Cerasicoccus_maritimus | 0 | 89.5 |
| Cereibacter_changlensis | 0 | 100.0 |
| Cetobacterium_ceti | 0 | 97.6 |
| Chelatococcus_albus | 0 | 99.2 |
| Chitinibacter_tainanensis_DSM_15459 | 0 | 100.0 |
| Chitinilyticum_aquatile_DSM_21506 | 0 | 99.2 |
| Chitinivibrio_alkaliphilus_ACht1 | 0 | 88.7 |
| Chitinolyticbacter_albus | 0 | 99.2 |
| Chitinophaga_cymbidii | 0 | 97.6 |
| Chlamydia_abortus | 0 | 72.6 |
| Chloracidobacterium_validum | 0 | 84.7 |
| Chlorobaculum_thiosulfatiphilum | 0 | 95.2 |
| Chlorobium_phaeovibrioides | 1 | 94.4 |
| Chloroflexus_islandicus | 0 | 96.0 |
| Chloroherpeton_thalassium_ATCC_35110 | 0 | 96.0 |
| Chondromyces_apiculatus_DSM_436 | 0 | 95.2 |
| Christensenella_massiliensis | 0 | 98.4 |
| Chromatium_okenii | 0 | 95.2 |
| Chromobacterium_indicum | 0 | 98.4 |
| Chromohalobacter_marismortui | 0 | 99.2 |
| Chryseobacterium_aureum | 0 | 100.0 |
| Chryseolinea_lacunae | 0 | 97.6 |
| Chrysiogenes_arsenatis_DSM_11915 | 0 | 97.6 |
| Chthoniobacter_flavus | 0 | 87.9 |
| Citreicella_sp._C3M06 | 0 | 99.2 |
| Citreimonas_salinaria | 0 | 98.4 |

| Species | diazotrophy | BUSCO |
|---|---|---|
| Citricoccus_alkalitolerans | 0 | 100.0 |
| Citrobacter_amalonaticus | 0 | 100.0 |
| Citromicrobium_bathyomarinum_JL354 | 0 | 99.2 |
| Clavibacter_lycopersici | 0 | 94.4 |
| Cloacibacterium_rupense | 0 | 100.0 |
| Clostridioides_difficile | 0 | 98.4 |
| Cohaesibacter_gelatinilyticus | 0 | 100.0 |
| Cohnella_faecalis | 1 | 99.2 |
| Collinsella_acetigenes | 0 | 93.5 |
| Colwellia_asteriadis | 0 | 100.0 |
| Comamonas_aquatilis | 0 | 100.0 |
| Compostimonas_suwonensis | 0 | 97.6 |
| Conexibacter_arvalis | 0 | 96.8 |
| Congregibacter_litoralis_KT71 | 0 | 99.2 |
| Coprococcus_ammoniilyticus | 0 | 99.2 |
| Coprothermobacter_platensis_DSM_11748 | 0 | 83.9 |
| Coraliomargarita_parva | 0 | 88.7 |
| Corallococcus_caeni | 0 | 99.2 |
| Coriobacterium_glomerans_PW2 | 0 | 91.1 |
| Corynebacterium_accolens | 0 | 100.0 |
| Costertonia_aggregata | 0 | 98.4 |
| Couchioplanes_azureus | 0 | 98.4 |
| Craurococcus_roseus | 0 | 100.0 |
| Crenothrix_polyspora | 0 | 97.6 |
| Croceibacter_atlanticus_HTCC2559 | 0 | 100.0 |
| Croceitalea_marina | 0 | 100.0 |
| Crocinitomix_catalasitica_ATCC_23190 | 0 | 99.2 |
| Crocosphaera_chwakensis_CCY0110 | 1 | 97.6 |
| Cronobacter_condimenti_1330 | 0 | 97.6 |
| Crossiella_equi | 0 | 97.6 |
| Cryobacterium_inferilacus | 0 | 96.0 |
| Cryomorpha_ignava | 0 | 98.4 |
| Cryptosporangium_minutisporangium | 0 | 99.2 |
| Cucumibacter_marinus_DSM_18995 | 0 | 100.0 |
| Cupriavidus_consociatus | 0 | 100.0 |
| Curtobacterium_caseinilyticum | 0 | 98.4 |
| Cutibacterium_acnes | 0 | 99.2 |
| Cyanothece_sp._BG0011 | 1 | 93.5 |
| Cyclobacterium_plantarum | 0 | 98.4 |
| Cycloclasticus_pugetii_PS-1 | 0 | 100.0 |
| Cytophaga_aurantiaca_DSM_3654 | 0 | 96.8 |
| Dactylosporangium_roseum | 0 | 97.6 |
| Dechloromonas_denitrificans | 0 | 99.2 |
| Deferribacter_thermophilus | 0 | 96.8 |
| Deferrisoma_camini_S3R1 | 0 | 97.6 |
| Defluviitoga_tunisiensis | 0 | 87.1 |
| Dehalobacter_restrictus_DSM_9455 | 0 | 99.2 |
| Dehalococcoides_mccartyi_195 | 0 | 91.9 |
| Dehalogenimonas_alkenigignens | 0 | 87.1 |
| Deinococcus_depolymerans | 0 | 91.9 |
| Delftia_deserti | 0 | 100.0 |
| Demequina_subtropica | 0 | 100.0 |
| Denitratisoma_oestradiolicum | 0 | 98.4 |
| Denitrobacterium_detoxificans | 0 | 93.5 |
| Denitrovibrio_acetiphilus_DSM_12809 | 0 | 96.8 |
| Dermabacter_jinjuensis | 0 | 96.8 |
| Dermacoccus_profundi | 0 | 99.2 |
| Dermatophilus_congolensis | 0 | 100.0 |
| Derxia_gummosa_DSM_723 | 1 | 99.2 |
| Desemzia_incerta | 0 | 94.4 |
| Desulfacinum_hydrothermale_DSM_13146 | 0 | 96.0 |
| Desulfarculus_baarsii_DSM_2075 | 0 | 97.6 |
| Desulfatiglans_anilini_DSM_4660 | 0 | 91.9 |
| Desulfatirhabdium_butyrativorans_DSM_18734 | 0 | 92.7 |
| Desulfatitalea_alkaliphila | 0 | 94.4 |
| Desulfitispora_alkaliphila | 0 | 100.0 |
| Desulfitobacterium_chlororespirans_DSM_11544 | 1 | 100.0 |

| Species | diazotrophy | BUSCO |
|---|---|---|
| Desulfobacca_acetoxidans_DSM_11109 | 0 | 95.2 |
| Desulfobacter_hydrogenophilus | 0 | 93.5 |
| Desulfobacterium_sp._N47 | 0 | 91.9 |
| Desulfobacula_phenolica | 0 | 93.5 |
| Desulfobaculum_xiamenense | 0 | 94.4 |
| Desulfobotulus_alkaliphilus | 0 | 94.4 |
| Desulfobulbus_elongatus_DSM_2908 | 0 | 85.5 |
| Desulfocapsa_sulfexigens_DSM_10523 | 0 | 96.0 |
| Desulfococcus_multivorans | 0 | 96.0 |
| Desulfocurvus_vexinensis_DSM_17965 | 0 | 94.4 |
| Desulfofustis_glycolicus_DSM_9705 | 0 | 94.4 |
| Desulfohalobium_retbaense_DSM_5692 | 0 | 96.8 |
| Desulfomicrobium_macestii | 0 | 94.4 |
| Desulfomonile_tiedjei_DSM_6799 | 1 | 96.0 |
| Desulfonatronospira_thiodismutans_ASO3-1 | 0 | 96.0 |
| Ensifer_oleiphilus | 1 | 100.0 |
| Enterobacter_adelaidei | 1 | 99.2 |
| Gluconacetobacter_liquefaciens | 1 | 100.0 |
| Heliobacterium_chlorum | 1 | 100.0 |
| Heliorestis_acidaminivorans | 1 | 99.2 |
| Herbaspirillum_lusitanum | 1 | 98.4 |
| Lamprobacter_modestohalophilus | 1 | 100.0 |
| Leptospirillum_ferriphilum | 1 | 83.1 |
| Magnetococcus_marinus_MC-1 | 1 | 99.2 |
| Magnetospira_sp._QH-2 | 1 | 100.0 |
| Magnetospirillum_sulfuroxidans | 1 | 99.2 |
| Mangrovibacterium_lignilyticum | 1 | 99.2 |
| Marichromatium_gracile | 1 | 99.2 |
| Marinobacterium_sediminicola | 1 | 100.0 |
| Mesorhizobium_calcicola | 1 | 96.0 |
| Methanococcus_voltae_PS | 1 | 17.7 |
| Methanosarcina_baikalica | 1 | 23.4 |
| Methanospirillum_stamsii | 1 | 21.8 |
| Methanothermobacter_defluvii | 1 | 21.0 |
| Methanothermococcus_thermolithotrophicus_DSM_2095 | 1 | 19.4 |
| Methylobacter_svalbardensis | 1 | 99.2 |
| Methylocapsa_aurea | 1 | 100.0 |
| Methylocella_tundrae | 1 | 98.4 |
| Methylocystis_silviterrae | 1 | 100.0 |
| Methyloglobulus_morosus_KoM1 | 1 | 99.2 |
| Methylomonas_defluvii | 1 | 98.4 |
| Methylovirgula_ligni | 1 | 100.0 |
| Moorella_naiadis__nom._illeg._ | 1 | 99.2 |
| Niveispirillum_irakense_DSM_11586 | 1 | 100.0 |
| Nostoc_piscinale_CENA21 | 1 | 91.9 |
| Orenia_metallireducens | 1 | 100.0 |
| Paenibacillus_alvei | 1 | 100.0 |
| Phaeospirillum_tilakii | 1 | 99.2 |
| Pleomorphomonas_diazotrophica | 1 | 100.0 |
| Pontibacter_locisalis | 1 | 98.4 |
| Propionispira_raffinosivorans_DSM_20765 | 1 | 99.2 |
| Rhizobium_alarense | 1 | 99.2 |
| Rhodobacter_lacus | 1 | 99.2 |
| Rhodocyclus_purpureus | 1 | 99.2 |
| Rhodoferax_potami | 1 | 99.2 |
| Rhodopila_globiformis | 1 | 96.8 |
| Rhodopseudomonas_telluris | 1 | 100.0 |
| Rhodospirillum_centenum_SW | 1 | 100.0 |
| Rubrivivax_gelatinosus | 1 | 99.2 |
| Sinorhizobium_arboris_LMG_14919 | 1 | 100.0 |
| Sphingomonas_aquatica | 1 | 99.2 |
| Telmatospirillum_siberiense | 1 | 100.0 |
| Thiococcus_pfennigii | 1 | 99.2 |
| Thiocystis_minor | 1 | 100.0 |
| Thiorhodococcus_fuscus | 1 | 100.0 |
| Tistlia_consotensis | 1 | 98.4 |
| Treponema_berlinense | 1 | 80.6 |

| **Species** | **diazotrophy** | **BUSCO** |
|---|---|---|
| Trichodesmium_erythraeum_IMS101 | 1 | 97.6 |
| Xanthobacter_flavus | 1 | 100.0 |

#### Supplementary Table S8

Complete plant species list for the nodulation analysis, including nodulation status and genome-quality metadata used in the OrthoGLMM model.

| Species | nodulation | BUSCO |
|---|---|---|
| Acacia_melanoxylon | 1 | 58.3 |
| Acmispon_strigosus | 1 | 60.0 |
| Adenanthera_pavonina | 0 | 62.4 |
| Aeschynomene_evenia | 1 | 60.3 |
| Albizia_julibrissin | 1 | 60.0 |
| Ammopiptanthus_mongolicus | 1 | 53.7 |
| Amphicarpaea_edgeworthii | 1 | 62.9 |
| Anthyllis_vulneraria | 1 | 63.0 |
| Apios_priceana | 1 | 56.2 |
| Arabidopsis_thaliana | 0 | 65.2 |
| Arachis_hypogaea | 1 | 64.6 |
| Astragalus_alpinus | 1 | 61.8 |
| Bauhinia_variegata | 0 | 62.9 |
| Cajanus_cajan | 1 | 58.5 |
| Calophaca_sinica | 1 | 57.4 |
| Canavalia_gladiata | 1 | 59.9 |
| Cannabis_sativa | 0 | 61.8 |
| Caragana_arborescens | 1 | 57.4 |
| Castanospermum_australe | 0 | 57.5 |
| Ceratonia_siliqua | 0 | 57.2 |
| Cercis_chuniana | 0 | 62.1 |
| Chamaecrista_fasciculata | 1 | 62.0 |
| Cicer_arietinum | 1 | 61.0 |
| Clitoria_ternatea | 1 | 56.5 |
| Coffea_arabica | 0 | 65.1 |
| Crotalaria_pallida | 1 | 61.7 |
| Cucumis_sativus | 0 | 59.6 |
| Cyamopsis_tetragonoloba | 1 | 62.3 |
| Dalbergia_candenatensis | 1 | 58.9 |
| Delonix_regia | 0 | 60.0 |
| Dicorynia_guianensis | 1 | 59.4 |
| Dipteryx_alata | 0 | 8.7 |
| Ebenus_cretica | 1 | 60.4 |
| Eperua_oleifera | 0 | 53.3 |
| Erythrophleum_ivorense | 1 | 68.6 |
| Eucalyptus_grandis | 0 | 60.0 |
| Flemingia_macrophylla | 1 | 57.4 |
| Fragaria_vesca | 0 | 63.4 |
| Gastrolobium_bilobum | 1 | 56.9 |
| Genista_pilosa | 1 | 59.2 |
| Gleditsia_sinensis | 0 | 58.2 |
| Glycine_max | 1 | 62.5 |
| Glycyrrhiza_glabra | 1 | 62.3 |
| Helianthus_annuus | 0 | 62.2 |
| Hoita_strobilina | 1 | 57.3 |
| Hylodesmum_podocarpum | 1 | 62.2 |
| Lablab_purpureus | 1 | 60.5 |
| Laburnum_anagyroides | 1 | 62.1 |
| Lathyrus_oleraceus | 1 | 61.0 |
| Lespedeza_cuneata | 1 | 25.2 |
| Leucaena_leucocephala | 1 | 65.8 |
| Lotus_japonicus | 1 | 61.5 |
| Lupinus_angustifolius | 1 | 63.1 |
| Macrotyloma_geocarpum | 1 | 61.4 |
| Malus_sylvestris | 0 | 63.0 |
| Medicago_truncatula | 1 | 63.9 |
| Mimosa_bimucronata | 1 | 61.2 |
| Mucuna_pruriens | 1 | 58.6 |
| Nissolia_brasiliensis | 0 | 57.9 |
| Ononis_reclinata | 1 | 66.4 |
| Ornithopus_perpusillus | 1 | 61.3 |
| Oxytropis_ochrocephala | 1 | 55.5 |
| Pachyrhizus_erosus | 1 | 57.9 |

| **Species** | **nodulation** | **BUSCO** |
|---|---|---|
| Phaseolus_vulgaris | 1 | 61.5 |
| Populus_trichocarpa | 0 | 63.9 |
| Prunus_persica | 0 | 63.9 |
| Psophocarpus_tetragonolobus | 1 | 57.2 |
| Pueraria_montana_var._lobata | 1 | 60.1 |
| Pyrus_communis | 0 | 62.8 |
| Retama_dasycarpa | 1 | 51.3 |
| Rosa_chinensis | 0 | 63.9 |
| Saraca_asoca | 0 | 46.0 |
| Senna_tora | 0 | 61.5 |
| Sesbania_bispinosa | 1 | 57.7 |
| Sindora_glabra | 0 | 58.1 |
| Solanum_lycopersicum | 0 | 60.3 |
| Sophora_flavescens | 1 | 51.8 |
| Spatholobus_suberectus | 1 | 56.8 |
| Sphenostylis_stenocarpa | 1 | 58.1 |
| Strongylodon_macrobotrys | 1 | 60.7 |
| Stylosanthes_angustifolia | 1 | 63.4 |
| Styphnolobium_japonicum | 0 | 58.5 |
| Theobroma_cacao | 0 | 63.1 |
| Trigonella_corniculata | 1 | 61.6 |
| Ulex_minor | 1 | 61.8 |
| Vachellia_pachyceras | 1 | 56.8 |
| Vicia_villosa | 1 | 62.8 |
| Vigna_angularis | 1 | 57.4 |
| Vitis_vinifera | 0 | 57.6 |

#### Supplementary Table S9

Complete mammalian species list for the marine lifestyle analysis, including phenotype labels and genome-quality metadata used in the OrthoGLMM model.

| Species | marine | BUSCO |
|---|---|---|
| Ailuropoda_melanoleuca | 0 | 93.3 |
| Alces_alces | 0 | 92.5 |
| Antilocapra_americana | 0 | 94.7 |
| Arctocephalus_gazella | 1 | 95.7 |
| Arctocephalus_townsendi | 1 | 92.1 |
| Balaenoptera_acutorostrata | 1 | 94.5 |
| Balaenoptera_bonaerensis | 1 | 50.4 |
| Balaenoptera_borealis | 1 | 94.7 |
| Balaenoptera_edeni | 1 | 94.8 |
| Balaenoptera_musculus | 1 | 93.2 |
| Balaenoptera_physalus | 1 | 94.7 |
| Bos_grunniens | 0 | 85.7 |
| Bos_taurus | 0 | 94.9 |
| Bubalus_bubalis | 0 | 94.8 |
| Callithrix_jacchus | 0 | 94.9 |
| Callorhinus_ursinus | 1 | 95.0 |
| Camelus_bactrianus | 0 | 96.1 |
| Camelus_dromedarius | 0 | 94.7 |
| Canis_latrans | 0 | 91.8 |
| Canis_lupus_baileyi | 0 | 95.9 |
| Caperea_marginata | 1 | 61.0 |
| Capra_hircus | 0 | 94.8 |
| Castor_canadensis | 0 | 95.4 |
| Cavia_porcellus | 0 | 95.0 |
| Ceratotherium_simum_simum | 0 | 94.8 |
| Cervus_elaphus | 0 | 95.4 |
| Choloepus_hoffmanni | 0 | 84.4 |
| Cnephaeus_nilssonii | 0 | 98.0 |
| Dasypus_novemcinctus | 0 | 93.6 |
| Delphinapterus_leucas | 1 | 94.2 |
| Delphinus_delphis | 1 | 94.3 |
| Desmodus_rotundus | 0 | 97.5 |
| Dugong_dugon | 1 | 94.3 |
| Elephas_maximus_indicus | 0 | 95.3 |
| Enhydra_lutris_kenyoni | 1 | 95.4 |
| Eptesicus_fuscus | 0 | 96.2 |
| Equus_asinus | 0 | 95.9 |
| Equus_caballus | 0 | 96.0 |
| Erignathus_barbatus | 1 | 90.9 |
| Erinaceus_europaeus | 0 | 93.0 |
| Eschrichtius_robustus | 1 | 94.9 |
| Eubalaena_glacialis | 1 | 95.4 |
| Eubalaena_japonica | 1 | 71.5 |
| Eumetopias_jubatus | 1 | 95.1 |
| Felis_catus | 0 | 96.1 |
| Giraffa_camelopardalis_rothschildi | 0 | 92.0 |
| Globicephala_melas | 1 | 94.5 |
| Gorilla_gorilla_gorilla | 0 | 95.8 |
| Grampus_griseus | 1 | 94.7 |
| Halichoerus_grypus | 1 | 95.6 |
| Hippopotamus_amphibius_kiboko | 0 | 95.4 |
| Homo_sapiens | 0 | 95.7 |
| Hydrurga_leptonyx | 1 | 95.9 |
| Hyperoodon_ampullatus | 1 | 93.8 |
| Ictidomys_tridecemlineatus | 0 | 95.3 |
| Inia_geoffrensis | 1 | 94.2 |
| Kogia_breviceps | 1 | 93.6 |
| Lagenorhynchus_albirostris | 1 | 94.3 |
| Leptonychotes_weddellii | 1 | 77.0 |
| Lepus_europaeus | 0 | 95.6 |
| Leucopleurus_acutus | 1 | 94.5 |
| Lipotes_vexillifer | 1 | 91.7 |
| Loxodonta_africana | 0 | 94.9 |

| Species | marine | BUSCO |
|---|---|---|
| Macaca_mulatta | 0 | 95.8 |
| Marmota_monax | 0 | 95.4 |
| Mesocricetus_auratus | 0 | 95.0 |
| Mesoplodon_bidens | 1 | 96.9 |
| Miniopterus_schreibersii | 0 | 98.8 |
| Monodelphis_domestica | 0 | 88.2 |
| Myotis_daubentonii | 0 | 98.4 |
| Myotis_lucifugus | 0 | 82.7 |
| Myotis_nattereri | 0 | 98.8 |
| Myrmecophaga_tridactyla | 0 | 63.3 |
| Notamacropus_eugenii | 0 | 90.5 |
| Ochotona_princeps | 0 | 95.0 |
| Odocoileus_virginianus | 0 | 93.3 |
| Orcinus_orca | 1 | 97.6 |
| Orycteropus_afer_afer | 0 | 90.8 |
| Oryctolagus_cuniculus | 0 | 96.0 |
| Ovis_aries | 0 | 95.2 |
| Pan_troglodytes | 0 | 95.5 |
| Panthera_leo | 0 | 93.7 |
| Panthera_tigris | 0 | 95.9 |
| Papio_anubis | 0 | 93.3 |
| Peromyscus_maniculatus_bairdii | 0 | 95.8 |
| Phascolarctos_cinereus | 0 | 86.8 |
| Phocoena_phocoena | 1 | 97.4 |
| Phyllostomus_discolor | 0 | 94.6 |
| Physeter_macrocephalus | 1 | 94.2 |
| Plecotus_auritus | 0 | 98.4 |
| Pongo_abelii | 0 | 95.7 |
| Procavia_capensis | 0 | 90.0 |
| Procyon_lotor | 0 | 80.9 |
| Pteropus_vampyrus | 0 | 86.9 |
| Rattus_norvegicus | 0 | 95.4 |
| Rattus_rattus | 0 | 91.6 |
| Rhinoceros_unicornis | 0 | 92.5 |
| Rhinolophus_ferrumequinum | 0 | 97.1 |
| Rhinolophus_hipposideros | 0 | 98.4 |
| Rhinolophus_sinicus | 0 | 96.4 |
| Saccopteryx_bilineata | 0 | 97.9 |
| Saccopteryx_leptura | 0 | 97.8 |
| Sagmatias_obliquidens | 1 | 94.2 |
| Sarcophilus_harrisii | 0 | 87.8 |
| Sciurus_carolinensis | 0 | 92.0 |
| Tursiops_truncatus | 1 | 97.3 |
